# Meningeal IL-17 producing T cells mediate cognitive impairment in salt-sensitive hypertension

**DOI:** 10.1101/2022.09.05.506398

**Authors:** Monica M. Santisteban, Samantha Schaeffer, Antoine Anfray, Giuseppe Faraco, David Brea Lopez, Gang Wang, Melissa Sobanko, Rose Sciortino, Gianfranco Racchumi, Ari Waisman, Laibaik Park, Josef Anrather, Costantino Iadecola

## Abstract

Hypertension, a disease afflicting over one billion individuals worldwide, is a leading cause of cognitive impairment, the mechanisms of which remain poorly understood. In a mouse model of hypertension, we found that the neurovascular and cognitive dysfunction depends on IL-17, a cytokine elevated in hypertensive individuals. However, neither circulating IL-17 or brain angiotensin signaling could account in full for the dysfunction. Rather, IL-17 produced by T-cells in the dura mater was the major culprit by reaching the cerebrospinal fluid and activating IL-17 receptors on brain associated macrophages. Accordingly, depleting brain macrophages, deleting IL17-RA in brain macrophages, or suppressing meningeal T cells completely rescued cognitive function without attenuating blood pressure elevation, circulating IL-17 or brain angiotensin signaling. The data unveil a critical role of meningeal T-cells and macrophage IL-17 signaling in the neurovascular and cognitive dysfunction of hypertension and suggest novel therapies to counteract the devastating effects of hypertension on cognitive health.

## INTRODUCTION

Hypertension (HTN) is a major cause of death and disability worldwide, and a leading risk factor for dementia^1^. Although there have been significant advances in the pharmacotherapy, a sizable proportion of patients have uncontrolled or resistant HTN which is particularly damaging to the brain^2, 3^. Furthermore, despite suggestive evidence that a rigorous control of blood pressure (BP) may lower the risk of mild cognitive impairment^4^, the burden of HTN on the brain remains substantial, including a 10% risk of recurrent cerebrovascular events despite BP control and no proven strategy to prevent dementia^5^. Therefore, there is need to gain a deeper understanding the damaging effects of HTN on the brain and to develop new approaches to protect cognitive health. Dysfunction of vital cerebrovascular regulatory mechanisms, such as the ability of neural activity to adjust the delivery of cerebral blood flow (CBF; functional hyperemia) or the regulation of microvascular perfusion by endothelial cells, have been strongly implicated in the deleterious effects of HTN on the brain^6^. However, the cellular and molecular basis through which the factors involved in BP elevation drive the neurovascular dysfunction associated with cognitive impairment remain poorly understood.

Salt-sensitivity is a critical factor in essential HTN^7^, affecting approximately 50% of hypertensive individuals^8^. Experimental studies using the deoxycorticosterone acetate (DOCA)-salt model have provided evidence that the renin-angiotensin system (RAS) is activated in brain and suppressed in the periphery^9, 10^. Indeed, a large proportion of individuals with resistant HTN, particularly African American and women, exhibit low levels of circulating renin, a key protease needed for angiotensin II (Ang II) production, suggesting suppression of systemic RAS^11, 12^. It is also well established that HTN induces immune dysregulation and elevates circulating levels of the cytokine interleukin (IL)-17 both in animals and humans^13–15^. Interestingly, high dietary salt increases circulating levels of IL-17 by promoting polarization of T-helper 17 lymphocytes (Th17) in the gut and induces neurovascular dysfunction and cognitive impairment^16–18^. However, the role IL-17 in the deleterious effects of salt-sensitive HTN on cognitive function, its sources and targets, and its relationships with brain RAS remain unexplored.

Here, we used the DOCA-salt model to examine the role of IL-17 in the neurovascular and cognitive dysfunction associated with salt-sensitive HTN. We found that DOCA-salt HTN alters key homeostatic mechanisms controlling the cerebral blood supply and leads to cognitive impairment. These deleterious effects are not driven by central Ang II signaling, but are associated with IL-17 signaling on both sides of the blood-brain barrier (BBB). In the circulation, IL-17 derived from gut and circulating T-cells activates IL-17 receptors A (IL-17RA) on cerebral endothelial cells to impair their ability to regulate cerebral perfusion, but this mechanism does not explain in full the cognitive deficits. On the brain side, unexpectedly, IL-17 derived from T-cells in the dura mater acts on IL-17RA on brain-associated macrophages (BAM)^19, 20^ to induce neurovascular uncoupling and cognitive impairment. Accordingly, depletion of meningeal T-cells or BAM completely rescues the cognitive phenotype. These findings unveil a previously unappreciated critical involvement of meningeal T-cells and IL-17 in the cognitive impairment associated with salt sensitive HTN and suggest novel approaches to ameliorate the deleterious impact of HTN on cognitive function.

## RESULTS

### Salt-sensitive HTN induces neurovascular and cognitive impairment linked to expansion of IL-17-producing cells

First, we sought to examine the impact of HTN on neurovascular and cognitive function. To this end, we used the DOCA-salt model of salt-sensitive HTN, in which mice are implanted with a s.c. pellet of DOCA and receive 0.9% NaCl in the drinking water^9, 21^. DOCA-salt treatment evoked a sustained elevation of BP beginning 3 days after pellet implantation (Fig 1A). An increase in circulating sodium was observed at 21 days (Suppl table 1), but the sodium content did not increase in brain, kidney and small intestine (Suppl Fig 1A). However, as previously reported in mouse models and in individuals with refractory HTN^22^, skin sodium content was increased without changes in potassium (Suppl Fig 1B). To examine the neurovascular effects of DOCA-salt HTN, we assessed CBF by laser-Doppler flowmetry in anesthetized mice with a cranial window overlying the somatosensory cortex under close monitoring of key physiological variables (Fig 1B; Methods)^23, 24^. DOCA-salt attenuated the increase in CBF evoked by neural activity induced by mechanical stimulation of the facial whiskers (functional hyperemia; Fig 1B-D), as well as the increase in CBF produced by bathing the somatosensory cortex with acetylcholine (ACh; Fig 1E), a response dependent on endothelial nitric oxide (NO)^25^. Both responses were impaired starting at day 10 after DOCA. However, smooth muscle vasoactivity, tested by neocortical application of adenosine (Fig 1F), BBB permeability to low molecular weight dextran (Suppl Fig 2A-B), and resting CBF (ml/100g/min) assessed by arterial spin label (ASL)-MRI (Suppl Fig 2C-D) were not impaired at day 21 after DOCA, indicating that the suppression of functional hyperemia and endothelial vasodilatation did not result from widespread neurovascular damage. DOCA-salt also altered cognitive function, as demonstrated by a reduction in the mice ability to discriminate between familiar and novel objects (working memory) (Fig 1G), a reduction of time spent in the target quadrant during the Barnes Maze probe trial (spatial learning and memory) (Fig 1H), and impaired nest building ability (activities of daily living) (Fig 1I). Representative images from each group are shown for each of these cognitive tasks (Fig 1G-I). Importantly, the neurovascular alterations preceded the development of cognitive impairment (Fig 1D, E, G), an observation consistent with a mechanistic link between neurovascular dysfunction and cognitive impairment^26^.

**Figure 1.**
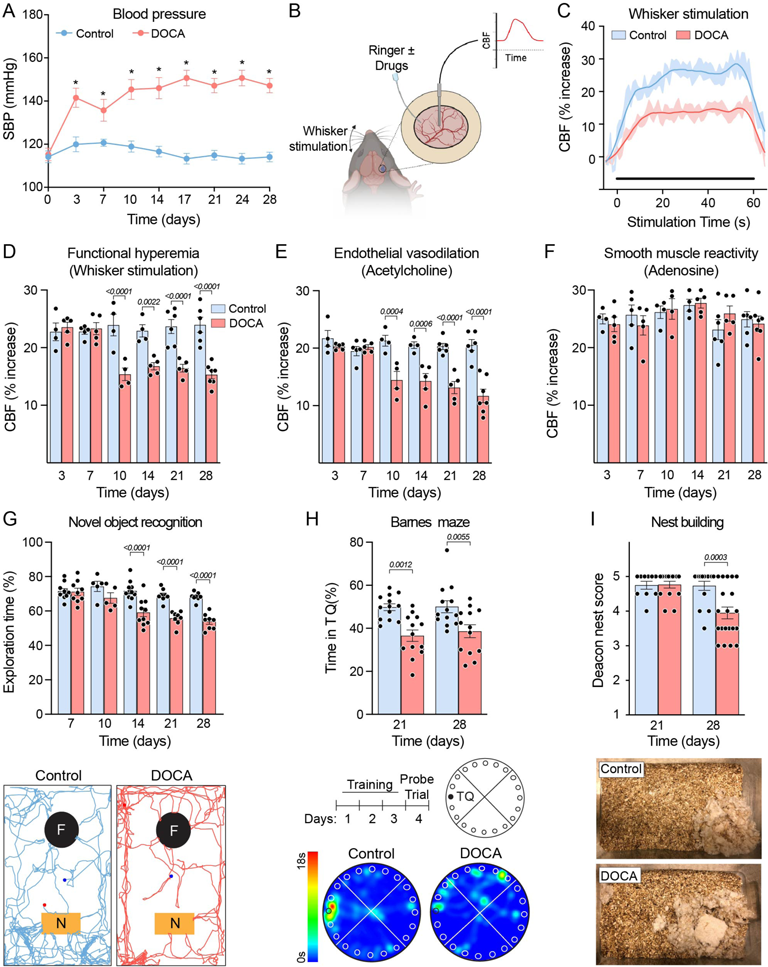
DOCA-salt hypertension induces neurovascular and cognitive impairment. **(A)** Systolic blood pressure (SBP), assessed by tail cuff plethysmography, is elevated in DOCA-salt hypertension over 28 days of treatment (HTN: p<0.0001, time: p<0.0001, interaction: p<0.0001; n=15). **(B)** Schematic of methods used to assess neurovascular function. **(C-D)** DOCA attenuates the increase in cerebral blood flow (CBF) induced by 60s stimulation of the facial whiskers (functional hyperemia), beginning at 10 days of DOCA (HTN: p<0.0001, time: p=0.001, interaction: p<0.0001; n=4-7). **(E)** Endothelial vasodilation was attenuated by DOCA beginning at 10 days (HTN: p<0.0001, time: p<0.0001, interaction: p<0.0001; n=4-7). **(F)** No difference was observed in smooth muscle reactivity (HTN: p=0.9702, time: p=.1981, interaction: p=.6479; n=4-7). **(G-I)** DOCA induced cognitive impairment assessed by **(G)** percent time exploring a novel object (HTN: p<0.0001, time: p<0.0001, interaction: 0.0012; n=5-11), **(H)** time in target quadrant (TQ) during Barnes maze probe trial (HTN: p<0.0001, time: p=0.6531, interaction: 0.7123; n=13), and **(I)** nest building assessed on the Deacon score scale (HTN: p=0.0125, time: p=0.0069, interaction: p=0093; n=10-20). Representative images of a single mouse from each group are shown for novel object recognition, Barnes maze probe trial, and nest building. All intergroup differences analyzed by two-way ANOVA and Bonferroni’s multiple comparison test. Data are shown as mean ± SEM.

Based on the emerging role of IL-17 in human HTN^13–15, 27, 28^ and in dietary salt-induced cognitive impairment^24^, we then examined whether IL-17 contributes to the neurovascular and cognitive effects of DOCA-salt HTN. Circulating IL-17 increased gradually over the course of the DOCA-salt treatment (Fig 2A), beginning at day 10 when neurovascular dysfunction first became apparent (Fig 1D-E). Focusing on the gut, an organ enriched with IL-17-producing cells^29^, we observed that 21 days of DOCA-salt increased *Il17a* mRNA expression (Fig 2B). To identify the cellular sources of IL-17, we induced DOCA-salt HTN in mice carrying the gene encoding eGFP at the *Il17a* locus^30^ and observed increased IL17-GFP+ cells at 21 days (Fig 2C-D) in the small intestine lamina propria, identified by flow cytometry to be Th17 and γδT17 cells (Fig 2E-G, Suppl Fig 3). Th17 and ψ8T17 cells have been shown to enter the circulation^30, 31^, and they were also increased in blood and spleen in DOCA-salt (Fig 2H-I), as in patients with HTN in whom circulating IL17-producing cells are increased^27, 28^. We found no evidence of IL-17 production by neutrophils (Suppl. Fig 4).

**Figure 2.**
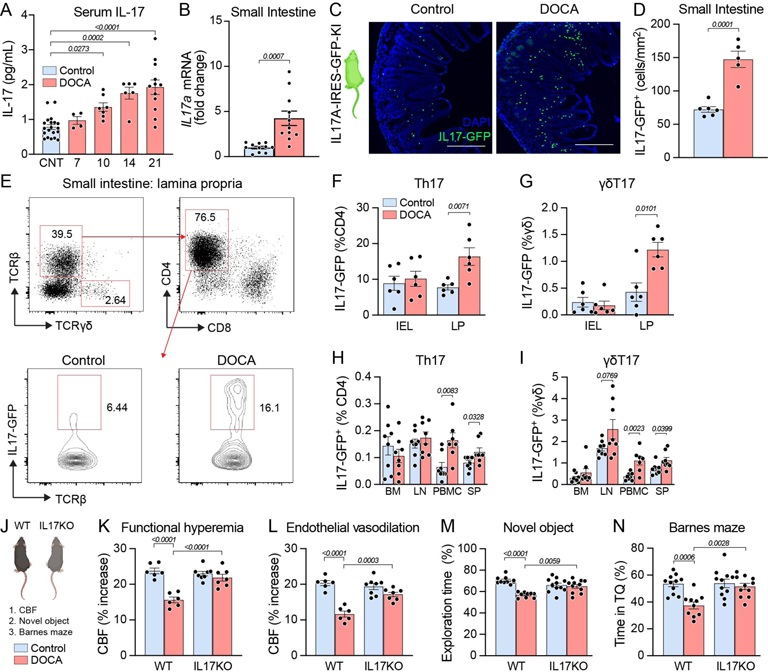
The neurovascular and cognitive impairment induced by DOCA is mediated by IL-17. **(A)** Serum IL-17 is elevated in DOCA HTN starting at 10 days (p<0.0001; one-way ANOVA and Bonferroni’s multiple comparison test; n=5-18 as shown). **(B)** 21 days of DOCA-salt increased *Il17a* mRNA (unpaired two-tailed t-test, n=11) as well as **(C-D)** IL17-GFP cells in the small intestine. (unpaired two-tailed t-test); Scale bar: 300μm. **(E)** By flow cytometry, these cells were identified to be **(F)** Th17 and **(G)** ψ8T17 in the lamina propria (LP) not intraepithelial lymphocytes (IEL) (unpaired two-tailed t-test, n=6). **(H-I)** Th17 and ψ8T17 were also expanded in peripheral blood mononuclear cells (PBMC) and spleen (SP), but not in the bone marrow (BM) or lymph nodes (LN) (unpaired two-tailed t-test per organ, n=7-8). **(J-N)** IL-17 deficient mice (KO) did not exhibit an attenuation in functional hyperemia (HTN: p<0.0001, genotype: p=0.0024, interaction: p=0.0002; two-way ANOVA and Bonferroni’s multiple comparison test; n=6-8) and endothelial vasodilation (HTN: p<0.0001, genotype: p=0.0079, interaction: p=0.0004; two-way ANOVA and Bonferroni’s multiple comparison test; n=6-8), and no deficits were observed in either novel object (HTN: p<0.0001, genotype: p=2603, interaction: p=0.0003; two-way ANOVA and Bonferroni’s multiple comparison test; n=9-11) or Barnes maze tests (HTN: p=0.001, genotype: p=0.0076, interaction: p=0.0114; two-way ANOVA and Bonferroni’s multiple comparison test; n=9-11). Data are shown as mean ± SEM.

To test whether IL-17 contributes to the deleterious effects of salt-sensitive HTN, we induced DOCA-salt HTN in IL-17 deficient mice (IL17KO; Fig 2J, Suppl table 2). IL17KO mice developed an increase in BP and circulating sodium similar to wild-type (WT) mice (Suppl Fig 5A, Suppl Table 1), but did not exhibit an attenuation in functional hyperemia and endothelial vasodilation (Fig 2K-L). Furthermore, no deficits were observed in either novel object or Barnes maze tests (Fig 2M-N). Thus, IL-17 produced by Th17 and γδT17 cells is essential for the neurovascular and cognitive dysfunction in DOCA-salt HTN.

### IL-17 impairs endothelial vasodilation by downregulating NO bioavailability via endothelial IL-17 receptors

Next, we sought to identify the cellular targets of the IL-17 contributing to neurovascular and cognitive impairment. Since endothelial cells are in direct contact with circulating IL-17, which is elevated in DOCA salt HTN (Fig 2A), we first assessed the contribution of cerebral endothelial IL-17RA. To this end, we deleted brain endothelial IL-17RA by administering an adeno-associated virus expressing Cre recombinase in cerebral endothelial cells (AAV-BR1-iCre; i.v.)^32, 33^ to IL-17RA^flox/flox^ mice^34^, referred to as IL-17RA^bECKO^. AAV-BR1-iCre delivery in Ai14-ROSA^tdTomato^ reporter mice (Suppl Fig 6A) demonstrated widespread endothelial cell transduction in the cerebral microvasculature (Suppl Fig 6B). Most importantly, we observed 90-95% endothelial viral transduction in vessels less than 20μm (Suppl Fig 6C-D), which include the arterioles involved in CBF regulation^35^. Three weeks after AAV-BR1-iCre delivery in IL-17RA^flox/flox^, we observed a reduction in *Il17ra* genomic DNA in sorted brain endothelial cells but not in microglia (Supp Fig 6E-F), consistent with the selectivity of this viral vector^33, 36^. IL-17RA^bECKO^ mice (Fig 3A) had increases in BP and circulating IL-17 comparable to those of DOCA-salt WT mice (Suppl Fig 5B, Suppl table 2), but the CBF response to ACh was completely rescued (Fig 3B). However, no improvement was observed in functional hyperemia (Fig 3C). Since the IL-17 has been shown to suppress endothelial NO production by inducing inhibitory eNOS phosphorylation at Thr^495 24^, we also examined NO production and endothelial nitric oxide synthase (eNOS) phosphorylation in DOCA-salt treated mice. Resting and ACh-induced endothelial NO production was attenuated in DOCA cerebral microvascular preparations (Fig 3D), an effect associated with an increase in eNOS inhibitory phosphorylation (Fig 3E). However, as predicted by the rescue of endothelial vasoactivity (Fig 3B), the increase in eNOS Thr^495^ phosphorylation was suppressed in IL-17RA^bECKO^ DOCA-salt mice (Fig 3F), attesting to the link between endothelial IL-17RA and eNOS inhibitory phosphorylation. Consistent with the partial rescue in neurovascular function, IL-17RA^bECKO^ DOCA-salt mice displayed cognitive improvement only at novel object recognition, not the Barnes maze (Fig 3G-H).

**Figure 3.**
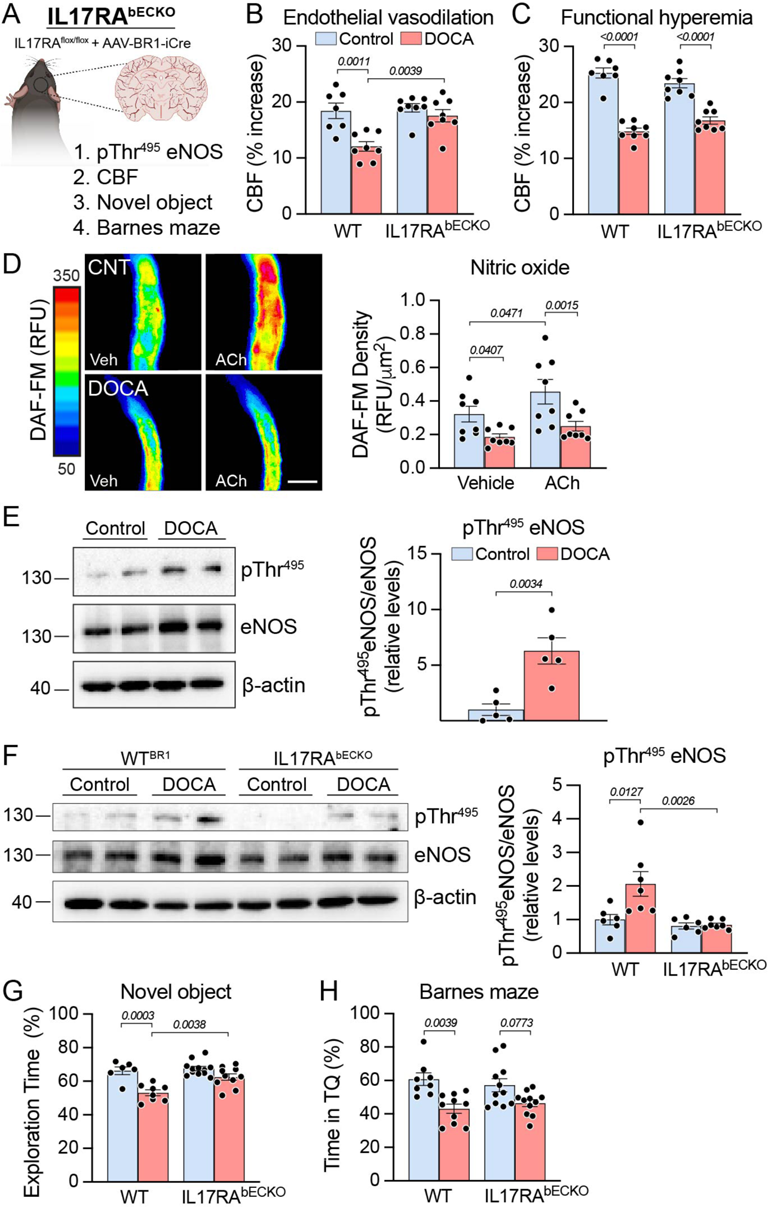
IL-17 impairs endothelial vasodilation by downregulating NO bioavailability via endothelial IL-17 receptors. **(A-C)** IL17RA brain endothelial cell knockout (IL17RAb^ECKO^) mice are protected from the impairment in endothelial vasodilation (HTN: p=0.0007, genotype: p=0.0067, interaction: p=0.0223; two-way ANOVA and Bonferroni’s multiple comparison test; n=7-8) but not the impairment in functional hyperemia induced by DOCA (HTN: p<0.0001, genotype: p=0.9446, interaction: p=0.0161; two-way ANOVA and Bonferroni’s multiple comparison test; n=7-8). **(D)** Resting and ACh-induced endothelial NO production was attenuated in DOCA cerebral microvascular preparations (p<0.0001, repeated measured one-way ANOVA and Tukey’s multiple comparison test; n=8). Scale bar: 50μm. **(E-F)** eNOS inhibitory phosphorylation was increased by DOCA in WT mice (unpaired two-tailed t-test), an effect suppressed in IL-17RAb^ECKO^ (HTN: p=0.0185, genotype: p=0.0037, interaction: p=0.0266; two-way ANOVA and Bonferroni’s multiple comparison test; n=5-7). **(G-H)** IL17RAb^ECKO^ displayed cognitive improvement only at novel object recognition (HTN: p<0.0001, genotype: p=0.0061, interaction: p=0.0338; two-way ANOVA and Tukey’s multiple comparison test; n=6-12), not the Barnes maze (HTN: p<0.0001, genotype: p=0.9599, interaction: p=0.2875; two-way ANOVA and Tukey’s multiple comparison test; n=6-12). Data are shown as mean ± SEM.

### BAM contribute to the neurovascular and cognitive dysfunction in DOCA-salt HTN

Cerebral endothelial IL-17RA knockdown ameliorated the neurovascular and cognitive function only partially, while total IL17KO rescued the dysfunction in full, suggesting the involvement of IL-17RA on other vessel-associated cell types. BAM, including perivascular and leptomeningeal macrophages, express IL-17RA^37^ and have been implicated in models of neurovascular and cognitive dysfunction^23, 38^, raising the possibility that they may also play a role DOCA-salt HTN. To test this hypothesis, we examined the effect of BAM depletion via intracerebroventricular (i.c.v.) delivery of liposome-encapsulated clodronate (Fig 4A)^23, 38^. Liposomes containing vehicle (PBS) or clodronate were injected i.c.v. on the same day as DOCA-salt treatment was started. This protocol depleted 80% of perivascular and leptomeningeal BAM (Fig 4B-C) within the time-frame of the experiment^23^, without affecting microglia or blood leukocytes^23^, BBB permeability^33^, or neurovascular function^23, 38^. Dural macrophages were depleted initially but, unlike the leptomeningeal and perivascular BAM, were fully restored within 21 days (Suppl Fig 7A-B). The BP (Suppl Fig 5C) and serum IL-17 (Suppl table 2) increases evoked by DOCA-salt were not affected by clodronate treatment. However, BAM depletion completely normalized functional hyperemia (Fig 4D), and partially improved endothelial vasoactivity (Fig 4E). Additionally, BAM depletion improved cognitive function as assessed by novel object recognition and Barnes maze (Fig 4F-G). These observations demonstrate that BAM contribute to the deleterious neurovascular and cognitive effects of DOCA-salt HTN.

**Figure 4.**
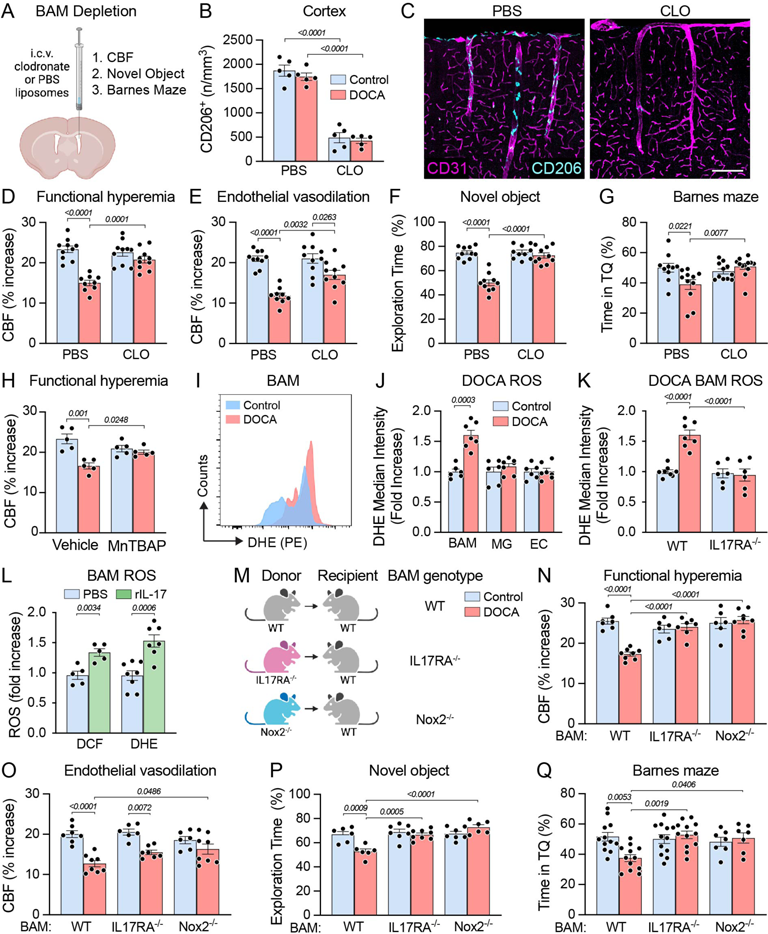
IL-17 impairs functional hyperemia via enhanced free radical production mediated by IL-17RA in BAM. **(A-C)** Brain associated macrophage (BAM), including perivascular and leptomeningeal macrophages, were depleted by 80% 21 days after intracerebroventricular delivery of liposome-encapsulated clodronate depletes (HTN: p=0.3156, liposomes: p=<0.0001, interaction: p=0.7304; two-way ANOVA and Tukey’s multiple comparison test; n=5). Scale bar: 150μm. **(D-G)** BAM depletion normalized functional hyperemia (HTN: p<0.0001, liposomes: p=0.0050, interaction: p=0.0004; two-way ANOVA and Tukey’s multiple comparison test; n=9-10) and partially improved endothelial vasoactivity (HTN: p<0.0001, liposomes: p=0.01, interaction: p=0.0112, two-way ANOVA and Tukey’s multiple comparison test; n=9-10) while also improving cognitive function assessed by novel object recognition (HTN: p<0.0001, liposomes: p<0.0001, interaction: p<0.0001, two-way ANOVA and Bonferroni’s multiple comparison test; n=10) and Barnes maze (HTN: p=0.1240, liposomes: p=0.0648, interaction: p=0.0062, two-way ANOVA and Bonferroni’s multiple comparison test; n=10-12). **(H)** Neocortical application of the ROS scavenger MnTBAP rescued the impairment of functional hyperemia in DOCA-salt (p=0.0017, repeated measures one-way ANOVA with Tukey’s multiple comparison test, n=5). **(I-K)** The increased ROS production in BAM induced by DOCA-salt (HTN: p=0.0033, cell type: p<0.0001, interaction: p<0.0001; two-way repeated measures ANOVA with Bonferroni’s multiple comparisons test; n=6-7) is prevented in IL-17RA deficient mice (HTN: p=0.0006, genotype: p<0.0001, interaction: p=0.0002; two-way ANOVA with Bonferroni’s multiple comparison test; n=6-8). **(L)** Recombinant IL-17 increased ROS production in BAM (unpaired two-tailed t-test per ROS indicator, n=5-8). **(M-Q)** Deletion of either IL-17RA or Nox2 from BAM in BM chimeras prevented the impairment of functional hyperemia in full (HTN: p=0.0030, BAM genotype: p=0.0003, interaction: p<0.0001; two-way ANOVA with Tukey’s multiple comparison test; n=6-8), improved endothelial vasoactivity (HTN: p<0.0001, BAM genotype: p=0.1898, interaction: p=0.0248; two-way ANOVA with Tukey’s multiple comparison test; n=6-8), as well as cognitive function (novel object HTN: p=0.0392, BAM genotype: p<0.0001; interaction: p=0.0004; Barnes HTN: p=0.2287, BAM genotype: p=0.0362, interaction: p=0.0040; two-way ANOVA with Tukey’s multiple comparison test; n=6-8). Data are shown as mean ± SEM.

### BAM contribute to neurovascular and cognitive dysfunction through IL17RA-dependent ROS production

Due to their myeloid origin, BAM are enriched with the ROS producing enzyme Nox2 and are major source of vascular oxidative stress^23, 39, 40^. Neocortical application of the ROS scavenger MnTBAP rescued the impairment of functional hyperemia in DOCA-salt (Fig 4H), attesting to the involvement of ROS in the neurovascular dysfunction. Therefore, we sought to determine if DOCA-salt increases ROS production in BAM and, based on the results in IL17KO mice (Fig 2J-N), whether the effect is IL-17 dependent. Dissociated brain cells from WT control and DOCA-salt mice were incubated with the ROS probe dihydroethidium (DHE; Fig 4I) and stained for identification of BAM (CD45^hi^CD11b^+^CD36^+^; Suppl Fig 8A,D)^23, 41^, microglia (CD45^int^CD11b^+^)^33, 42^, and endothelial cells (CD45^-^Ly6C^+^)^33, 42^ using flow cytometry^23, 33, 41, 42^. We found that DOCA-salt increased ROS production in BAM, but not in microglia or endothelial cells (Fig 4J). DOCA-salt failed to increase ROS in BAM of IL-17RA deficient mice (Fig 4K), indicating that IL-17 signaling is needed for BAM ROS production in DOCA-salt HTN. Consistent with this conclusion, recombinant IL-17 (10ng/mL) increased ROS production in BAM, as assessed with two different probes (Fig 4L).

To provide further evidence that IL-17RA in BAM are involved, we used a bone marrow (BM) chimera-based approach. We and others have demonstrated that BM transplantation after total body irradiation repopulates leptomeningeal and perivascular compartments with BM-derived macrophages^23, 33, 43, 44^. Therefore, we transplanted IL-17RA^-/-^ or Nox2^-/-^ BM into WT mice to replace BAM with IL-17RA^-/-^ or Nox2^-/-^ BM-derived cells (Fig 4M). Three months later, mice were placed on the DOCA-salt protocol. WT mice transplanted with WT BM (WT**→**WT) exhibited alterations in CBF responses and cognition identical to those observed in naïve mice (Fig 4N-Q, Fig 1D-H), indicating that although BAM in these mice are derived from the BM, they are pathogenically equivalent to native yolk sac-derived BAM^23, 38^. Deletion of either IL-17RA or Nox2 in BAM prevented the impairment of functional hyperemia in full (Fig 4N) and partially improved endothelial vasoactivity (Fig 4O), as observed in the BAM depletion experiments (Fig 4D-E). Additionally, IL-17RA^-/-^**→** WT and Nox2^-/-^**→** WT chimeras showed improved cognitive function (Fig 4P-Q). Attesting to the requirement of IL-17RA in BAM, ROS production was blunted in BAM from IL-17RA^-/-^**→** WT DOCA-salt chimeras (Suppl Fig 9B). Since ROS are a well-known product of IL17 signaling^45^ these data also provide evidence of IL17RA activation in BAM.

To complement the BM chimera data implicating IL17RA in BAM, we developed a novel mouse expressing tamoxifen-inducible Cre recombinase under the control of the Mrc1 promoter (Mrc1^CreERT2/+^; Fig. 5A). First, we sought to determine whether Mrc1^CreERT2^ heterozygosity altered BAM number and distribution across brain compartments (Suppl Fig 10A). To this end, after labeling BAM by i.c.v. injection of FITC dextran (Suppl Fig 10B), we confirmed that the number of BAM in dural, pial, and perivascular spaces of Mrc1^CreERT2/+^ mice were not different from those in Mrc1^+/+^ (WT) mice (Suppl Fig 10C-G). Next, to characterize the efficiency of this novel Cre driver mouse, we crossed Mrc1^CreERT2^ with Ai14 TdTM (Fig 5B-C). Following tamoxifen treatment, we found widespread TdTM expression in pial and perivascular BAM (Suppl Fig 11A-F). TdTM was not detected in Iba1+ microglia (Suppl Fig 11E). Tamoxifen treatment of adult Mrc1^CreERT2/+^ crossed with IL17-RA^flox/flox^ (BAM^IL17RA-/-^) led to a 78.9% reduction in IL17-RA+ BAM (Fig 5D-E, Suppl Fig 11G-H), without affecting the BP response (Fig 5F) and circulating IL-17 elevation (Suppl Table 2) induced by DOCA-salt. Consistent with the findings in IL-17RA^-/-^**→** WT chimeras, we observed full rescue of functional hyperemia without improvement of endothelial vasodilation (Fig. 5G-H). Additionally, BAM^IL17RA-/-^ DOCA-salt mice showed improved cognitive function (Fig 5I-J). Collectively, the findings with BM chimera, and our novel Mrc1^CreERT2/+^ mouse provide converging evidence that IL-17RA in BAM are critical for the alterations in functional hyperemia and cognitive function induced by DOCA-salt HTN.

**Figure 5.**
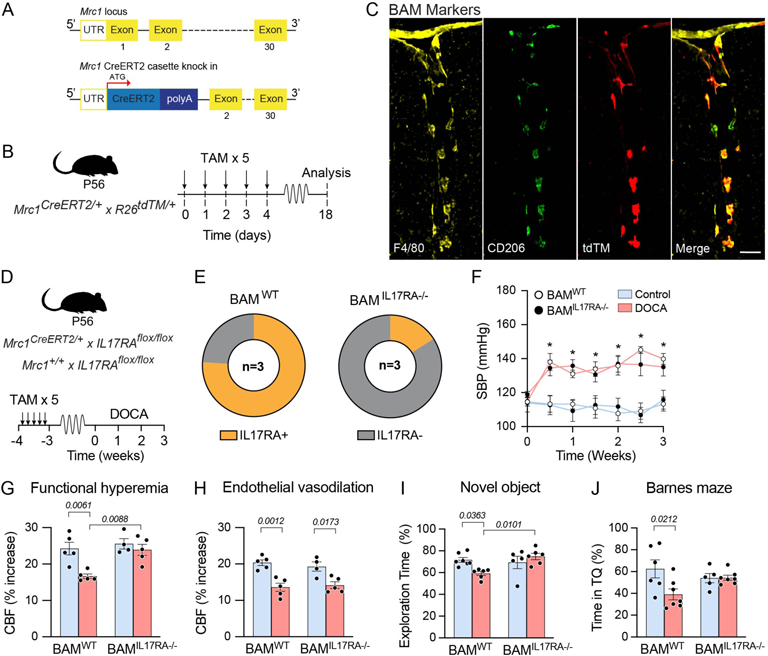
IL17-RA deletion in BAM improves functional hyperemia and cognitive function in DOCA-salt. **(A)** CreERT2 cassette was introduced into the endogenous Mrc1 locus. **(B)** These mice were crossed with the Ai14 TdTomato reporter mice, and **(C)** showed TdTM expression in BAM. Scale bar: 50μm. **(D)** Mrc1^CreERT2^ mice were crossed with IL17-RA^flox/flox^ to generate BAM^IL17RA-/-^. **(E)** IL17RA expression was reduced by 78.9% in BAM^IL17RA-/-^ compared to BAM^WT^ mice. (n=3 mice/group; 50 BAM per mouse; p<0.0001, two-way ANOVA with Bonferroni’s multiple comparisons test). **(F)** Blood pressure under control and DOCA-salt conditions was not altered in BAM^WT^ or BAM^IL17RA-/-^ mice (two-way repeated measures ANOVA with Tukey’s multiple comparisons test). **(F-G)** IL17-RA deletion in BAM restored functional hyperemia (HTN: p=0.0048, genotype: p=0.0080, interaction: p=0.0499; two-way ANOVA with Bonferroni’s multiple comparison test; n=4-5) without improving endothelial vasodilation (HTN: p<0.001, genotype: p=0.8120, interaction: p=0.4368; two-way ANOVA with Bonferroni’s multiple comparison test; n=4-5), and **(H-I)** rescued cognitive function after 21 days of DOCA-salt (novel object HTN: p=0.2638, genotype: p=0.0460; interaction: p=0.0100; Barnes HTN: p=0.0427, genotype: p=0.5297, interaction: p=0.0365; two-way ANOVA with Tukey’s multiple comparison test; n=5-7). Data are shown as mean ± SEM.

### DOCA-salt HTN increases IL-17-producing T cells in the dura mater

Next, we sought to define the cellular source(s) of IL-17 acting on BAM IL-17RA to induce neurovascular and cognitive dysfunction. Recent evidence indicates that IL-17-producing T cells are present in the dura mater^46, 47^, and are able to modulate rodent behavior^48, 49^. In agreement with these findings, *Il17a* mRNA was detected in the dura of control mice and was markedly increased by 21 days of DOCA-salt treatment (Fig 6A). *Il17a* mRNA was not observed in the brain parenchyma (Fig 6A). To map IL-17 producing cells in brain and dura we used IL17-GFP reporter mice. Consistent with the mRNA data, IL17-GFP+ cells were not observed in the brain (Suppl Fig 12), but were found in the dura (Fig 6B-D), as previously reported^48, 50^. DOCA-salt treatment (21 days) led to a significant increase in IL17-GFP+ cells surrounding the venous sinuses but not in regions removed from the sinus (Fig 6B-C). To determine whether these cells actually secrete IL-17, we performed an IL-17 ELISpot assay to detect cytokine release with single-cell resolution^51^ in isolated dural leukocytes. We found that dural cells secrete IL-17, and this response is increased in DOCA-salt mice (Fig 6E-F). We then used flow cytometry to characterize the IL17-GFP+ cells. We did not observe an increase of CD45+ immune cells in the dura of DOCA-salt mice (Fig 6G). We found no change in the total number of CD4 or Th17, (Suppl Fig 13A-B), or the percent of CD4 producing IL-17 (Fig 6H), excluding a role for these cells. There was no change in the total number of total ψ8T cells (Suppl Fig 13C) and a trend towards increase of ψ8T17 cells (Suppl Fig 13D). Interestingly, we observed an increase in the percentage of ψ8T cells producing IL17 (Fig 6I). Since the total number of ψ8T-cells did not change (Suppl Fig 13C), changes in other cells populations would not impact the % of ψ8Tcells producing IL17. Therefore, consistent with data in the literature^48^, we observed an increased in the proportion of dural ψ8T cells producing IL17. The increase in dural IL-17 production after 21 days of DOCA-salt was associated with increased detection of IL-17 in the cerebrospinal fluid (CSF; Fig 6J). To gain insight on how dura-derived IL-17 reaches the CSF, we assessed the structural integrity of the arachnoid barrier using immunocytochemistry for the arachnoid barrier tight junction marker claudin-11^52^. We found significant tight junction remodeling in DOCA-salt mice as evidenced by (1) discontinuous tight junctions (Fig 6K), (2) reduced arachnoid domain areas (Fig 6L-M), and (3) altered tight junction morphology (Fig 6K)^53, 54^. Collectively, these findings suggest that IL-17 generated by T-cells in the dura reach the CSF in the subarachnoid space via a disrupted arachnoid barrier and engage IL-17RA on BAM. Of note, we did not observe changes in key inflammatory genes (Suppl Fig 14A-B) or evidence of microglia and astrocyte activation (Suppl Fig 14C-D), suggesting that dura-derived IL17 did not result in a massive neuroinflammatory reaction which could potentially contribute to cognitive impairment in DOCA-salt.

**Figure 6.**
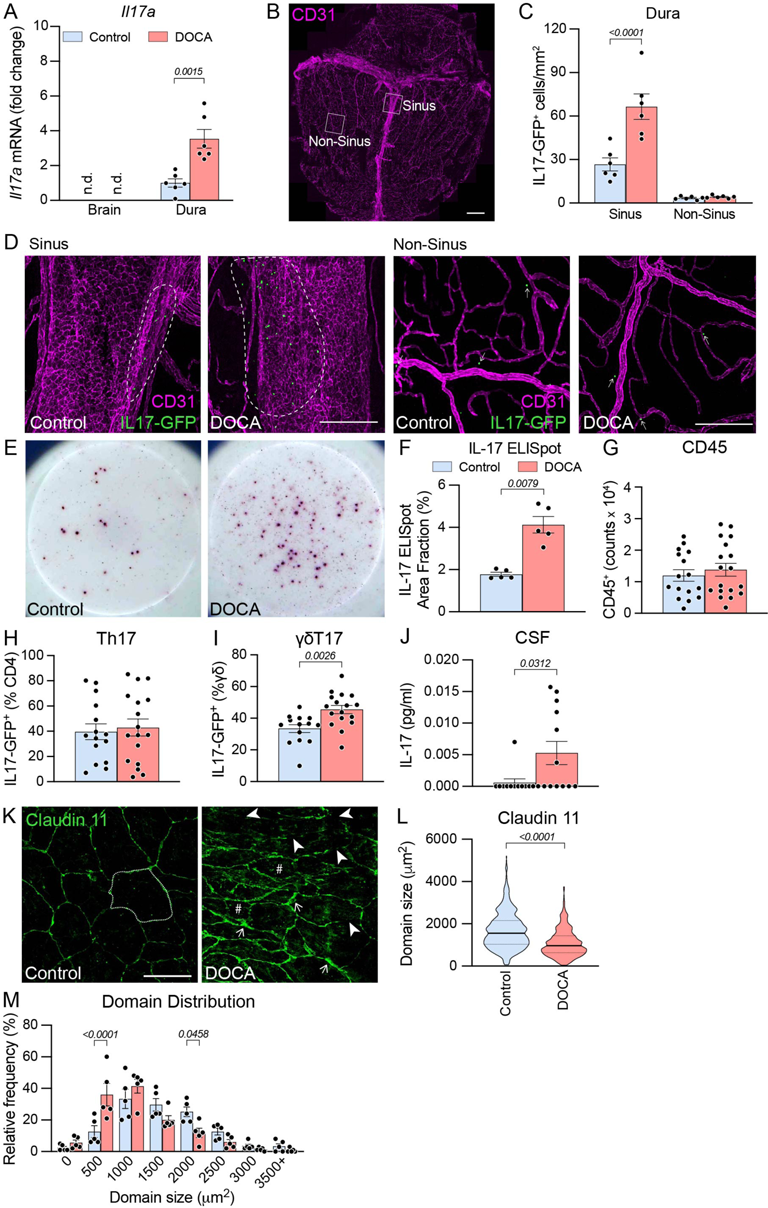
Salt-sensitive hypertension increases IL17-producing T cells located in the dura mater. **(A)** *Il17a* mRNA was not observed in the brain parenchyma, but it was detected in dura mater of control mice and was markedly increased by DOCA-salt (unpaired two-tailed t-test; n=6). **(B)** Dura whole mount stained with CD31 (magenta). Scale bar: 1mm. **(C)** DOCA-salt treatment led to a significant increase in IL17-GFP+ cells in the vicinity of the venous sinuses (HTN: p=0.0017, Sinus: p<0.0001, interaction: p=0.0034; two-way repeated measures ANOVA with Bonferroni’s multiple comparison test; n=6). **(D)** Representative images of control and DOCA sinus and non-sinus regions. Analysis shown in 6C. **(E-F)** Dural isolated cells secrete IL-17, and this response is increased in DOCA-salt mice (unpaired two-tailed t-test; n=5). **(G)** DOCA-salt does not change the total number of CD45 immune cells isolated from dura. (**H-I)** DOCA-salt increases the percentage of ψ8T17 cells but no difference in Th17 cells (unpaired two-tailed t-test; n=14-17 mice). **(J)** IL-17 detection was increased in the cerebrospinal fluid (CSF) after 21 days of DOCA-salt (Wilcoxon signed-rank test, n=12-13). **(K)** Arachnoid barrier integrity was assessed by the arachnoid barrier tight junction marker Claudin-11. Arrowheads indicate discontinuous tight junctions, # indicates reduced arachnoid domain size, and thin arrows indicate altered tight junction morphology. (**L-M**) Arachnoid barrier domain size (Kolmogrov-Smirnov test; n=616-663 domains [5 mice, 10 images per mouse, all domains per image quantified]) and distribution was altered by DOCA-salt (HTN: p=0.9880, domain size bin: p<0.0001; interaction: p<0.0001; two-way ANOVA with Bonferroni’s multiple comparison test). Data are shown as mean ± SEM.

### Cognitive impairment in salt-sensitive HTN is driven by dural IL17-producing ψ8T cells

Next, we sought to provide evidence in support of the involvement of dural IL17-producing cells in the neurovascular and cognitive effects of DOCA salt HTN. Dural ψ8T cells, like other ψ8T cells^55^, are tissue resident, with only 1-2% being derived from the circulation under homeostasis^48^. However, in inflammatory conditions, an increased influx of γδT cells into lymph nodes and subsequent homing to inflamed tissues via the circulation has been shown^56, 57^.

Given that circulating Th17 and γδT17 cells are both elevated in DOCA-salt, it is conceivable that T cells migrate from the circulation to the dura. To test this hypothesis, we utilized FTY720 (fingolimod), a sphingosine-1-phosphate receptor (S1PR) modulator that depletes circulating lymphocytes, including ψ8T17 and Th17 cells, by preventing their egress from lymphoid tissues and gut, as well as other mechanisms^31, 58–61^. FTY720 (1mg/kg i.p. every 3 days^24^) was administered from day 7 through day 21 of DOCA-salt treatment (Fig 7A) and did not affect the development of HTN (Fig 7B). As expected, FTY720 significantly reduced circulating CD4^+^ T cells (Fig 7C) without affecting the elevation in circulating IL-17 in DOCA-salt mice (Fig 7D). FTY720 depleted IL17-GFP+ cells in the dura (Fig 7E), both Th17 cells (Fig 7F), and ψ8T17 cells (Fig 7G). Interestingly, FTY720 did not ameliorate the CBF response evoked from the endothelium (Fig 7H), attesting to its dependence on circulating IL-17 acting on cerebral endothelial IL-17RA and not BAM. The reduction of IL-17 producing T cells in the dura was associated with full rescue of functional hyperemia (Fig 7I) and improved of cognitive function (Fig 7J-K), as well as suppression of ROS production in BAM (Suppl Fig 9C).

**Figure 7.**
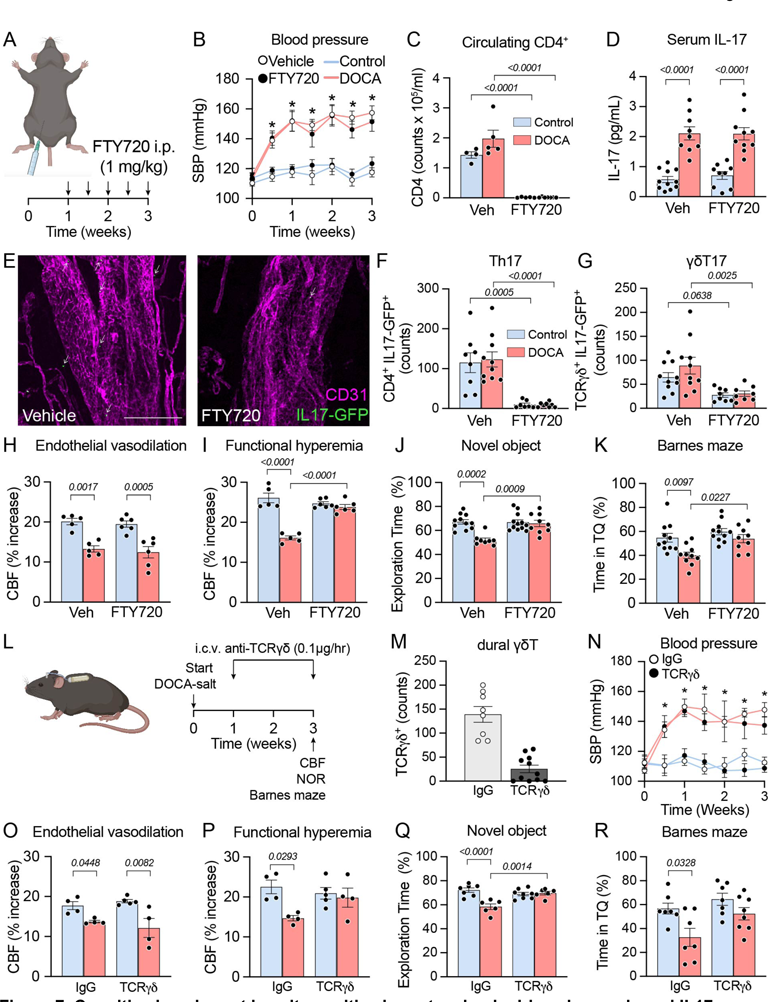
Cognitive impairment in salt-sensitive hypertension is driven by meningeal IL17-producing T cells. **(A-B)** FTY720 was administered from day 7 through day 21 of DOCA-salt and did not affect the increase in systolic blood pressure (treatment: p<0.0001, time: p<0.0001, interaction: p=0.0182; two-way repeated measures ANOVA with Tukey’s multiple comparison test; n=5-8). **(C-D)** FTY720 reduces circulating CD4 T cells (HTN: p=0.0705, treatment: p<0.0001, interaction: p=0.0737; two-way ANOVA with Tukey’s multiple comparison test; n=4-7), without affecting the elevation in serum IL-17 in DOCA-salt (HTN: p<0.0001, treatment: p=0.7200, interaction: p=0.6622; two-way ANOVA with Tukey’s multiple comparison test; n=9-11). **(E-G)** FTY720 reduced IL17-GFP cells in the meninges, including both Th17 (HTN: p=0.8358, treatment: p<0.0001, interaction: p=0.7848; two-way ANOVA with Bonferroni’s multiple comparison test; n=8-10) and ψ8T17 (HTN: p=0.2517, treatment: p=0.0003, interaction: p=0.3639; two-way ANOVA with Bonferroni’s multiple comparison test; n=8-10). **(H-K)** FTY did not improve endothelial vasoactivity (HTN: p<0.0001, treatment: p=0.4839, interaction: p=0.9293; two-way ANOVA with Bonferroni’s multiple comparison test; n=5-6), but completely restored functional hyperemia (HTN: p<0.0001, treatment: p=0.0004, interaction: p<0.0001; two-way ANOVA with Bonferroni’s multiple comparison test; n=5-6), as well as improved cognitive function (novel object HTN: p=0.0007, treatment: p=0.0037; interaction: p=0.0023; Barnes HTN: p=0.0023, treatment: p=0.0045, interaction: p=0.1695; two-way ANOVA with Tukey’s multiple comparison test; n=8-12). **(L)** anti-TCRgd antibody was delivered i.c.v. from day 7 to 21 of DOCA-salt. **(M)** anti-TCRgd antibody treatment reduced gdT cells in the dura (p<0.0001, two-tailed unpaired t-test). **(N)** anti-TCRgd antibody did not alter the blood pressure response to DOCA-salt (two-way repeated measures ANOVA with Tukey’s multiple comparisons test). **(O-R)** Dura gdT cell depletion did not restore endothelial vasodilation (HTN: p=0.0008, treatment: p=0.8541, interaction: p=0.2935; two-way ANOVA with Tukey’s multiple comparison test; n=4-5), but was associated with improved functional hyperemia (HTN: p=0.0190, treatment: p=0.3072, interaction: p=0.0655; two-way ANOVA with Tukey’s multiple comparison test; n=4-5) and cognitive function (novel object HTN: p=0.0012, treatment: p=0.0323; interaction: p=0.0004; Barnes HTN: p=0.0040, treatment: p=0.0229, interaction: p=0.3002; two-way ANOVA with Tukey’s multiple comparison test; n=6-8). Data are shown as mean ± SEM.

To confirm the role of dural ψ8T cells in the neurovascular and cognitive dysfunction in DOCA-salt, we infused anti-TCRψ8 antibody into the cerebral ventricles (i.c.v.) using osmotic minipumps from day 7 through day 21 of DOCA-salt treatment (Fig 7L) to deplete ψ8T cells in the dura (Fig 7M). i.c.v. delivery of this antibody has been previously shown to decrease ψ8T cell production of IL-17 in the dura^48^, but did not affect the BP response to DOCA-salt (Fig 7N). Consistent with results of the FTY720 experiments, ψ8T cell-depletion in the dura did not improve endothelial vasodilation (Fig 7O). The reduction of ψ8T cells in the dura rescued functional hyperemia (Fig 7P), and improved cognitive function (Fig 7Q-R). Thus, dural IL17-producing T-cells are the source of the IL-17 contributing to the neurovascular and cognitive impairment in salt-sensitive HTN.

### The contribution of Ang II to the cerebrovascular dysfunction in DOCA-salt depends on IL-17 signaling

It is well established that the DOCA-salt treatment leads to activation of brain RAS^9, 10^, which has been implicated in endothelial dysfunction in large and small cerebral vessels^62^. In apparent contrast, our data suggest that IL-17 is essential for the neurovascular and cognitive dysfunction in DOCA-salt HTN. Therefore, we investigated the relationship between brain Ang II and IL-17 in this model. As expected, DOCA-salt treatment elevated Ang II levels in brain and reduced it in the circulation (Fig 8A-B). These effects were associated with mRNA upregulation of brain Ang II receptors type 1A (AT1R) (*Agtr1a*; Fig 8C) and downregulation of kidney renin (*Ren1*; Fig 8D), confirming activation of RAS in brain and suppression in the periphery^62^. Central AT1R blockade by i.c.v. infusion of losartan (Fig 8E), prevented the increase in BP caused by DOCA-salt^9^ (Suppl Fig 5) but did not attenuate the increase in circulating IL-17 (Suppl table 2). Accordingly, i.c.v. losartan restored functional hyperemia (Fig 8F), but did not improve endothelium-dependent vasodilation (Fig 8G), consistent with circulating IL-17 being the predominant mediator of the endothelial dysfunction. Central AT1R blockade ameliorated cognitive impairment only partially, with an improvement observed only in novel object recognition, but not at the Barnes maze (Fig 8H-I). Since Ang II induces ROS production in BAM leading to neurovascular dysfunction^23^, we wondered if IL-17 signaling in BAM is required for Ang II-induced ROS production. Ang II stimulation increased ROS production in WT but not IL17RA^-/-^ BAM (Fig 8J). However, indices of brain RAS activation were not attenuated in IL17KO mice (Suppl Fig 15A) or mice in which dural production of IL-17 was suppressed by treatment with FTY720 (Suppl Fig 15B), suggesting that brain RAS activation in the absence of dural IL-17 is not sufficient to induce neurovascular dysfunction. In support of this conclusion, bathing the cerebral cortex with Ang II to activate AT1R on BAM^23^, induced neurovascular dysfunction in WT but not IL17KO mice (Fig 8K-L). Thus, IL-17 signaling is necessary for the contribution of Ang II to the neurovascular and cognitive impairment of salt-sensitive HTN.

**Figure 8.**
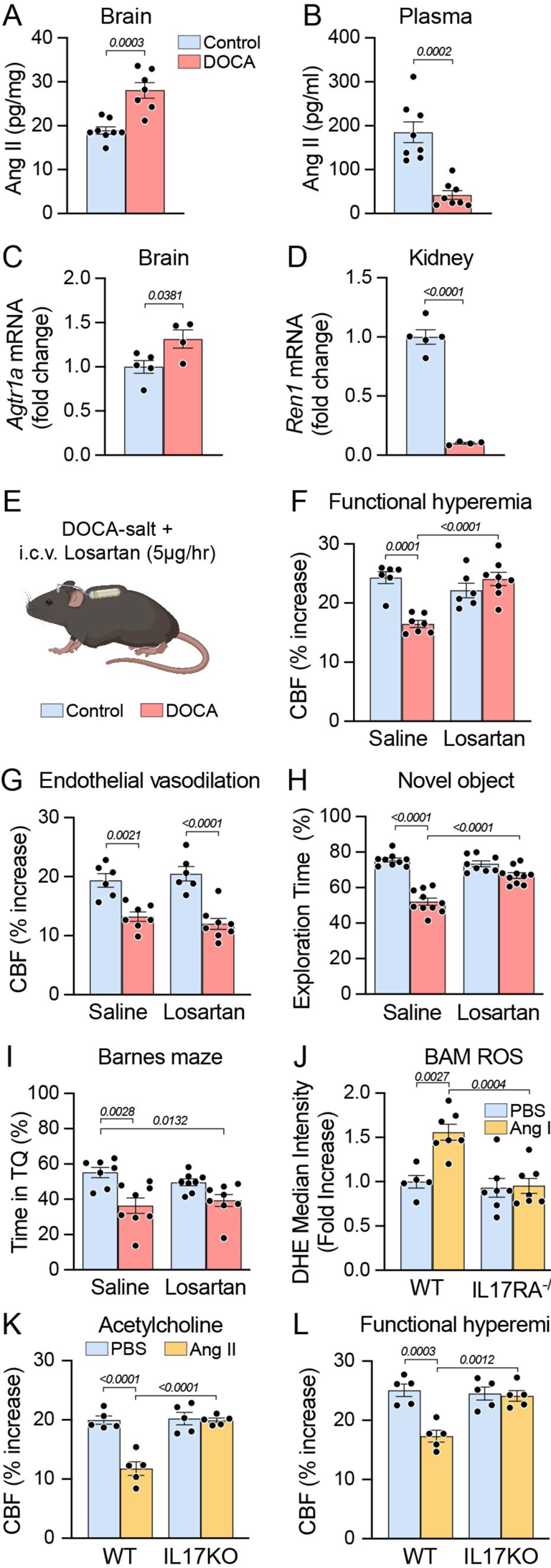
The contribution of Ang II to the cerebrovascular dysfunction in DOCA-salt depends on IL-17 signaling. **(A-B)** DOCA-salt treatment elevated Ang II levels in brain and reduced it in the circulation (unpaired two-tailed t-test; n=7-8). **(C-D)** DOCA-salt upregulates brain *Agtr1a* and downregulates kidney renin (unpaired two-tailed t-test; n=4-5). **(E-G)** Central AT1R blockade with i.c.v. losartan restored functional hyperemia (HTN: p=0.0088, treatment: p=0.0148, interaction: p<0.0001; two-way ANOVA with Bonferroni’s multiple comparison test; n=6-8), but did not improve endothelium-dependent vasodilation (HTN: p<0.0001, treatment: p=0.9604, interaction: p=0.2642; two-way ANOVA with Bonferroni’s multiple comparison test; n=6-8). **(H-I)** Central AT1R blockade improved novel object recognition (HTN: p<0.0001, treatment: p=0.0005, interaction: p<0.0001; two-way ANOVA with Tukey’s multiple comparison test; n=8-10), but did not improve Barnes maze (HTN: p=0.0002; treatment: p=0.6968, interaction: p=0.2113; two-way ANOVA with Tukey’s multiple comparison test; n=7-8). **(J)** Ang II stimulation increased ROS production in WT but not IL17RA^-/-^ BAM (genotype: p=0.0015; treatment: p=0.0050, interaction: p=0.0087; two-way ANOVA with Tukey’s multiple comparison test; n=5-7). **(K-L)** Neocortical application of Ang II impaired endothelial vasoactivity (genotype: p=0.0029; treatment: p=0.0003, interaction: p=0.0004; two-way repeated measures ANOVA with Bonferroni’s multiple comparison test; n=5) and induced neurovascular dysfunction (genotype: p=0.0374; treatment: p=0.0003, interaction: p=0.0005; two-way repeated measures ANOVA with Bonferroni’s multiple comparison test; n=5) in WT but not IL17KO mice. Data are shown as mean ± SEM.

## DISCUSSION

We have demonstrated that the neurovascular and cognitive dysfunction associated with salt-sensitive HTN is mediated by two distinct mechanisms involving IL-17 signaling in cerebral endothelium and BAM (Suppl Fig 16): (1) in the circulation, IL-17 produced by T-cells acts on cerebral endothelial IL-17RA to reduce NO production leading to suppression of endothelial vasoactivity without affecting the increase in CBF induced by neural activity; (2) in the brain, IL-17 produced by dura T-cells acts on IL-17RA on BAM to induce vascular oxidative stress and suppression of functional hyperemia with minimal effects on endothelial function. However, these two mechanisms do not contribute equally to the cognitive dysfunction: counteracting the effect of circulating IL-17 by endothelial IL-17RA deletion rescues cognition only partially, while counteracting the sources (T-cells) or targets of central IL-17 (BAM or BAM IL-17RA) rescues cognitive function in full. Furthermore, endothelial NO synthase phosphorylation is not involved in the mechanisms of neurovascular coupling since functional hyperemia can be rescued completely by BAM depletion or IL17RA deletion in BAM despite persistent endothelial vasomotor dysfunction. Thus, the deleterious effects of BAM on functional hyperemia cannot be attributed to endothelial dysfunction or eNOS phosphorylation. Therefore, the data unveils an unanticipated central role of meningeal T-cells in the deleterious cognitive effect of salt sensitive hypertension.

There is increasing evidence that inflammation and immunity participate in the pathobiology of HTN^63^. Pioneering studies have unveiled a role of innate and adaptive immunity in the central and peripheral mechanisms driving the elevation in BP and on the end organ damage, particularly in the kidney and the vasculature^64–66^. In this context, T-cells and IL-17 have emerged as important mediators of the effect of Ang II and DOCA-salt HTN on peripheral organs and vessels, in part verified in hypertensive patients^67^. Here, we extend these observations by providing evidence that dural T-cells and IL-17 play a critical role in the neurovascular and cognitive deficits in a model of HTN reproducing key attributes of the human disease^10, 68^. While the dura mater has recently been recognized as major players in the immune responses of the brain underling brain injury and repair^46, 48^, our findings provide insight into an previously unrecognized pathogenic role of meningeal immunity in HTN. Our studies revealed that the dura is the critical site of the immune responses underlying the neurovascular and cognitive deficits in HTN. This process is driven by a crosstalk between dural T-cells and BAM through IL-17 signaling. Thus, depletion of IL-17-producing T-cells in the dura, depletion of BAM or deletion of IL-17RA on BAM, rescues the cognitive deficits in full. As for the mechanisms by which BAM alter neurovascular coupling, evidence from our studies implicate Ang II^23^ and amyloid-beta^38^ inducing BAM production of ROS, which are thought to inactivate neuronal NO and other vasoactive mediators released from neurons and converging on resistance arteriole to mediate vasodilatation^35, 69^.

The DOCA salt model is associated with increases in brain RAS signaling and reduction of the systemic RAS^10^, which we have verified by central and peripheral measurements of Ang II. DOCA salt HTN is well established to induce alterations in cerebral vascular regulation both in vivo and in isolated cerebral arteries and arterioles^21, 62^. Surprisingly, however, our study showed that the vascular effects of Ang II, mediated by ROS production by BAM, require IL-17. Thus, ex vivo Ang II is unable to increase ROS production in BAM in the absence of IL-17RA, and in vivo Ang II applied to the neocortex to target meningeal and perivascular BAM does not induce neurovascular dysfunction IL17KO mice. These observations unveil a critical requirement of IL-17 in the deleterious effects of Ang II. Considering the key role that Ang II and IL-17 signaling play in health and disease the molecular basis of their interaction is of great interest and will require further exploration.

Owing to the absolute reliance of the brain on the delivery of blood flow, reduced cerebral perfusion or alterations in neurovascular regulation have long been implicated in cognitive impairment induced by vascular factors as well as Alzheimer’s disease^70, 71^. Our studies, in general, support a link between CBF regulation and cognitive health, but they also suggest effects of IL-17 independent of blood flow. While resting CBF is not reduced, blocking cerebral endothelial IL-17 signaling rescues endothelial vasoactivity and produces partial cognitive improvement. On the other hand, counteracting central Ang II signaling rescues only neurovascular coupling and leads to a partial improvement in cognition. Although it is conceivable that endothelial dysfunction and neurovascular uncoupling could play an additive role in the cognitive deficits, this possibility seems unlikely because depletion of dural T-cells provides complete cognitive rescue while improving only neurovascular coupling and not endothelial function. Direct effects of IL-17 on neurons have been demonstrated in other models^48, 49, 72, 73^, which in conjunction with the vascular effects could contribute to the cognitive dysfunction.

It is well established that HTN is a leading risk factor for cognitive impairment caused both by vascular factors and neurodegeneration^6^, but the evidence that antihypertensive therapy reduces such risk is inconsistent and limited^74, 75^. Our findings reveal an additional layer of complexity in the deleterious effects of HTN on the brain and suggest new preventive and therapeutic approaches. The central actions of Ang II are critical for the development of HTN by inducing neurohumoral dysfunction. This aspect was highlighted by our observation that central inhibition of AT1R prevented the BP elevation completely but did not rescue the cognitive dysfunction in full. While BP control remains critical for attenuating hypertensive end-organ damage to kidney, heart, and vasculature^76^, full protection of the brain may require also targeting meningeal immunity. Considering the diversity of mechanisms underlying human HTN^77^, efforts to select hypertensive patients in which immune factors are involved, may identify individuals at greater risk for the deleterious effects of dural immune signaling on the brain. Since the infectious complications of suppressing immune signaling is a well-known concern in patients with cardiovascular diseases^78^, strategies to selectively target dural immunity would be required^79^.

## MATERIALS AND METHODS

The data that support the findings of this study are available from the corresponding authors upon request. Details on antibodies used and specific primer sequences are found in the Online Supplement. For rigor and reproducibility, experiments were repeated at least twice throughout the manuscript, and figures present pooled data.

### Experimental mouse models

All procedures were approved by the Institutional Animal Care and Use Committee of Weill Cornell Medicine and performed in accordance with the National Institutes of Health (NIH) Guide for the Care and Use of Laboratory Animals. Studies were performed in a blinded fashion in male C57BL/6 mice (WT, age 3-5 months, weight 25-30g; JAX, the Jackson Laboratory), IL-17 GFP reporter mice (C57BL/6-*IL17a^tm1Bcgen^*/J, JAX strain# 018472), IL-17 knockout mice (*IL17a^tm^*^1^.^1^*^(icre)Stck^*/J, JAX strain# 016879), and IL-17RA^flox/flox^ mice^34^. IL-17RA^flox/flox^ mice were crossed with the germ-cell driven Sox2-Cre mice (B6.Cg-*Edil3^Tg(Sox^*^2^*^-cre)1Amc^*/J, JAX strain# 008454)^80^ to generate whole-body IL-17RA knockout mice. Bone marrow chimera experiments detailed below were performed on C57BL/6 mice receiving donor cells from IL-17RA knockout mice or B6.129S-*Cybb^tm1Din^*/J mice (*Nox2^−/−^*, JAX strain# 002365), both lines are congenic for C57BL/6. AAV-BR1-iCre experiments detailed below were performed on B6.Cg-*Gt(ROSA)26Sor^tm^*^14^*^(CAG-tdTomato)Hze^*/J (Ai14 TdTomato reporter, JAX strain# 007914), and homozygous IL-17RA^flox/flox^ mice. Mrc1^CreERT2^ mice were generated by Cyagen. The Mrc1 targeting construct was linearized by restriction digestion with NotI, followed by phenol/chloroform extraction and ethanol precipitation. The linearized vector was transfected into C57BL/6 ES cells according to Cyagen’s standard electroporation procedures. The transfected ES cells were subject to G418 selection (200 μg/mL) 24 hours post electroporation. 179 G418 resistant clones were picked and amplified in 96-well plates. Two copies of 96-well plates were made, one copy was frozen down and stored at −80°C and the other copy of the 96-well plates was used for DNA isolation and subsequence PCR screening for homologous recombination. The PCR screening identified 18 potential targeted clones, from among which 6 were expanded and further characterized by Southern blot analysis. Five of the six expanded clones were confirmed to be correctly targeted.

The Mrc1 gene (NCBI Reference Sequence: NM_008625.2) is located on mouse chromosome 2. Thirty exons have been identified, with the ATG start codon in exon 1 and the TAG stop codon in exon 30. In the targeting vector, the coding region of exon 1 plus part of intron 1 was replaced with the CreERT2-polyA cassette. Correct integration of the CreERT2-polyA cassette was confirmed by diagnostic PCR. Sequencing of the genomic region of interest after amplification by PCR was further used to verify correct integration. After germline transmission, correctly targeted founders were crossed with C57BL/6J mice to establish the colonies for this study. Adult 6-10 week old mice were treated with tamoxifen citrate (Sigma: T-5648; oral gavage; 200mg/kg) once a day for 5 consecutive days. Female mice were not included in this study, as previous studies have shown that they have a blunted hypertensive and immune response to DOCA-salt^81^.

### DOCA-salt HTN

Mice were randomized to treatment group and anesthetized by isoflurane inhalation for subcutaneous implantation of a 50 mg pellet of DOCA (Innovative Research of America, Cat# M-121) or sham surgery of equal duration^21^. After recovery from anesthesia, DOCA animals were maintained on standard chow and *ad libitum* access to 0.9% NaCl in autoclaved tap water. Control animals were maintained on standard chow and *ad libitum* access to autoclaved tap water. Systolic BP was monitored in awake mice using tail-cuff plethysmography (Hatteras)^23, 33^. Mice were acclimated to tail-cuff plethysmography for one week prior to pellet or sham surgery, and systolic BP was monitored twice a week after initiation of DOCA-salt.

### Tissue sodium measurement

Mice were anesthetized with isoflurane, and blood was collected through cardiac puncture. Brain, kidney, skin (dorsal abdomen), and distal small intestine (distal 10 cm) were isolated, flash frozen, and stored at −80°C until analysis. After 15 minutes, blood was centrifuged at 2,000*g* for 15 minutes and serum was separated and stored at −80°C until analysis. The serum renal chemical panel was performed by the Laboratory of Comparative Pathology (LCP) of Weill Cornell Medicine Research Animal Resource Center, and the tissue mineral panel for sodium measurement was performed by the Animal Health Diagnostic Center Toxicology Lab of Cornell University Veterinary Medicine using inductively coupled plasma – atomic emission spectrometry (ICP-AES)^82, 83^.

### General surgical procedures for CBF studies

Mice were anesthetized with isoflurane (induction, 5%; maintenance, 2%) and artificially ventilated with a mixture of N2 and O2. One of the femoral arteries was cannulated for recording mean arterial pressure (MAP) and collecting blood samples. Rectal temperature was maintained at 37°C. After surgery, isoflurane was discontinued and anesthesia was maintained with urethane (750 mg/kg, i.p.) and chloralose (50 mg/kg, i.p.). Throughout the experiment, the level of anesthesia was monitored by testing of motor responses to tail pinch. Arterial blood gases were monitored at the beginning and end of the experiment and maintained at pO2 100-110mmHg, pCO2 30-40mmHg, and pH 7.3-7.4^23^. As in previous studies^84^, MAP remained within the autoregulated range for CBF (Control: 82.77 ± 0.85 mmHg; DOCA: 106.37 ± 0.99 mmHg; p<0.05). Due to the anesthesia, the baseline BP and the increase in BP induced by DOCA-salt was lower than that observed in awake mice^9^.

### Experimental protocol for experiments monitoring CBF reactivity

As previously performed^23, 24, 38^, a small craniotomy (2 × 2 mm) was performed to expose the parietal cortex, the dura was removed, and the site was superfused with Ringer’s solution (37°C; pH 7.3–7.4)^23^. CBF was continuously monitored at the site of superfusion with a laser-Doppler probe (Perimed) positioned stereotaxically on the cortical surface and connected to a data acquisition system (PowerLab). CBF values were expressed as percentage increases relative to the resting level. After MAP and blood gases were stable, CBF responses were recorded. The whisker-barrel cortex was activated for 60 seconds by stroking of the contralateral vibrissae, and the evoked changes in CBF were recorded. ACh (10 μM; Sigma-Aldrich) or adenosine (400 μM; Sigma-Aldrich) was superfused on the exposed neocortex for 5 minutes^23, 24, 38^. In some experiments, CBF responses were tested before and after 30 minutes of superfusion with the ROS scavenger MnTBAP (100μM)^23, 84^ or Ang II (500nM)^23, 85^.

### BBB permeability

BBB permeability was assessed using fluorescein-dextran (FITC-dextran, MW 3kDa; 100μl of 1% solution i.v.), as previously described^23, 33^. The tracer was allowed to circulate for 20 minutes, and then mice were transcardially perfused with cold PBS to clear the intravascular tracer. Brains were removed and olfactory bulb, brainstem, and cerebellum were discarded. Samples were weighed and frozen on dry ice and stored at −80 °C until analysis. Tissue was homogenized in 400μL of PBS, mixed with 400μL of methanol, and centrifuged at 13,000*g* for 30 minutes. The supernatant was used for measurement of the amount of FITC-dextran (485nm excitation and 530 nm emission), measured in duplicate using a fluorescence spectrophotometer, together with standards, and normalized to brain tissue weight.

### Measurement of resting CBF by ASL-MRI

CBF was assessed quantitatively using arterial spin labeling magnetic resonance imaging (ASL-MRI) at 21 days of DOCA-salt, performed on a 7.0-tesla 70/30 Bruker Biospec small-animal MRI system with 450 mT/m gradient amplitude and a 4,500 T · m^−1^ · s^−1^ slew rate. A volume coil was used for transmission and a surface coil for reception. Anatomical localizer images were acquired to find the transversal slice approximately corresponding to bregma +0.5 mm. This position was used for subsequent ASL-MRI, which was based on a flow-sensitive alternating inversion recovery rapid acquisition with relaxation enhancement (FAIR-RARE) pulse sequence labeling the inflowing blood by global inversion of the equilibrium magnetization. One axial slice was acquired with a field of view of 15 × 15 mm, spatial resolution of 0.117 × 0.117 × 1 mm, TE (echo time) of 5.368 ms, effective TE of 48.32 ms, recovery time of 10 s, and a RARE (rapid imaging with refocused echoes) factor of 72. Twenty-two turbo inversion recovery values ranging from 30 to 2,300 ms were used, and the inversion slab thickness was 4 mm. For computation of resting CBF (rCBF), the Bruker ASL perfusion processing macro was used. It uses a published model^86^ and includes steps to mask out the background. The masked rCBF images were exported to Analyze format on the MRI console. The ASL images were analyzed by ImageJ and the average CBF value is reported in mL per 100 g of tissue per min^24^.

### Novel object recognition *test*

The novel object recognition test (NOR) task was conducted in a plastic box measuring 29 cm × 47 cm × 30 cm high^23, 24^. Stimuli consisted of plastic objects that varied in color and shape, but had similar size. A video camera was used to record the testing session for offline analysis using AnyMaze software. Mice were acclimated to the testing room for 1 hour each day prior to the start of each day. On day 1, mice were acclimated to the testing chamber (habituation). On day 2, mice were placed in the same chamber in the presence of 2 identical sample objects and were allowed to explore for 5 minutes. After an intersession interval of 1 hour, mice were placed in the same chamber, but 1 of the 2 objects was replaced by a novel object. Mice were allowed to explore for 5 minutes. Between trials, the maze is cleaned with 10% ethanol in water to minimize olfactory cues. Exploratory behavior was assessed manually by two experimenters blinded to the treatment group. Exploration of an object was defined as the mouse sniffing the object or touching the object while looking at it^23^. A minimal exploration time for both objects (total exploration time) during the test phase (5 seconds) was used. The amount of time taken to explore the novel object was expressed as percentage of the total exploration time and provides an index of working memory.

### Barnes Maze

As described previously^23, 24^, we used a Barnes maze consisting of a circular open surface (90 cm in diameter) elevated to 90cm by four wooden legs. There are 20 circular holes (5 cm in diameter) equally spaced around the perimeter, positioned 2.5cm from the edge of the maze. No wall or intra-maze visual cues are placed around the edge. A plastic escape box (11 x 6 x 5 cm) was positioned beneath one of the holes. Mouse movement is tracked with the Any-Maze software (Stoelting). Mice are tested in groups of 10, and between trials are placed into cages in a dark room adjacent to the test room for the intertrial interval (45 minutes). Mice are habituated to the dark room for 60 min prior to the start of each day. No habituation trial is performed. The acquisition phase consists of three consecutive training days with three 3-minute trials per day with the escape hole located at the same location across trials and days. On each trial, a mouse is placed into a start tube located in the center of the maze, the start tube is raised, and the buzzer is turned on until the mouse enters the escape hole. After each trial, mice remain in the escape box for 60s before being returned to their home cage. Between trials, the maze is cleaned with 10% ethanol in water to minimize olfactory cues. Three parameters of learning performance are recorded: (1) latency to locate the escape hole, (2) distance traveled before locating the escape hole, and (3) number of errors made. Errors are defined as head-pokes into non-escape holes and are counted manually. On the fourth day, the probe trial is performed and consists of a 1.5 min trial where the escape hole has been removed. The memory parameter recorded is percent of time spent in the target quadrant where the escape hole used to be.

### Nest building

The ability of mice to build nests is assessed by the Deacon rating scale^24, 87^. 1 hour prior to the dark cycle, each mouse is placed in a new clean cage containing 5g of nestlet (Ancare) in the middle of the cage. Food, water, and lighting parameters are not changed from standard housing practices. The next day, nests are assessed on a rating scale of 1-5^87^, and untorn nestlet pieces are weight. The cognitive parameters recorded are (1) nest score, and (2) percent of untorn nestlet.

### IL-17 measurement

IL-17 concentration in serum was measured by cytometric bead array mouse IL-17A Enhanced Sensitivity Flex Set (BDBiosciences)^24^ or by electrochemiluminescence-based multi-array MSD V-Plex Mouse IL-17A Kit (MesoScale)^88^, according to the manufacturer’s instructions.

### Immunofluorescence

IL17-eGFP and wild-type mice were anesthetized with sodium pentobarbital (120 mg/kg, i.p.) and perfused transcardially with phosphate-buffered saline (PBS) followed by 4% paraformaldehyde (PFA) in PBS. Distal small intestine, brain, and skull cap were removed and post-fixed overnight in 4% PFA. Small intestine was then submerged in 30% sucrose solution for 3 days, frozen, and sections (thickness: 30 µm) were cut through the whole distal small intestine using a cryostat and then place on a slide. Free-floating coronal brain sections (thickness 40 μm) were cut through the whole brain using a vibrating microtome. Dura whole mounts were prepared by careful stripping from the skull cap using fine surgical forceps, as has been previously described by us^24, 30, 89^ and others^48, 50, 90^. For leptomeningeal mounting, PFA-perfused brains were post-fixed in methanol on a shaker at room temperature for 10 minutes^52^. Then, the leptomeninges were carefully stripped from the surface of the brain, and mounted on slides for staining^52–54^. Brain sections and dura whole mounts were permeabilized in 0.5% Triton-PBS and then blocked with 5% of normal donkey or goat serum in 0.1% Triton-PBS. Sections were incubated with primary antibodies (Suppl Table 3) at 4°C overnight in 2% normal donkey or goat serum in 0.1% Triton-PBS. Leptomeningeal mounts were blocked with 5% normal donkey or goat serum and 5% BSA in 0.1% Triton-PBS. Sections were then incubated with species-specific secondary antibody (1:100, Jackson ImmunoResearch whole IgG affinity-purified antibodies) and mounted on slides with VectaShield Hardset mounting medium with DAPI (Vector Labs), visualized with a laser-scanning confocal microscope (Leica TCS SP8), and analyzed on ImageJ software by an investigator blinded to the treatment groups. IL17-GFP cells in the dura were analyzed as sinus-associated if they were in close proximity with the venous sinus, and non-sinus if they were in all other areas of the dura. For TdTM+ vessel quantification in AAV-BR1-Cre injected Ai14 TdTM reporter mice, 100 vessels were randomly selected from 10 different images of the cortex from 3 mice. The area of TdTM as a percent of CD31 immunofluorescence was quantified.

### Cell suspension preparation from lymph nodes, spleen, and blood

At the indicated timepoints, mesenteric, axillary and inguinal (non-brain draining) lymph nodes were extracted from each animal and pooled, placed on a premoistened 70-µm cell strainer, gently triturated, washed with 10 mL of PBS and spun at 500*g* for 7 min^24^. The cell suspension was then either stained for flow cytometry analysis or processed for analysis of intracellular cytokines. The spleen was removed, its epithelium was cut longitudinally, and cells were isolated as described for the lymph nodes. Blood (150 µL) was drained from the submandibular venous plexus into heparinized tubes, incubated with erythrocytes lysis buffer and spun at 500*g* for 7 min, and cells were stained for flow cytometry analysis^24^.

### Isolation of intestinal lamina propria mononuclear cells

Mice were euthanized by isoflurane overdose, and small intestines were removed and separated as previously described^24, 30^. Peyer patches were cut out from the small intestine and small intestines were completely cleaned of mesenteric fat and intestinal contents^24^. Then intestines were opened longitudinally, washed of fecal contents with PBS, cut into approximately 1 cm pieces and placed into 20 mL of HBSS/10 mM HEPES, 8% FBS, 4 mM EDTA, 0.5 mM DTT. Next intestinal pieces were washed three times in a shaking incubator set at 250 rpm and at 37 °C for 20 min. After each round, intestinal pieces were vortexed for 20 s and the cell suspension containing intraepithelial lymphocytes (IELs) was collected. Suspensions from the three washes of IELs were combined and filtered over 0.3 g of prewashed nylon wool placed into a 10-mL syringe and then over a 70-µm strainer. Intestinal pieces were washed with complete PBS to remove EDTA, minced thoroughly with scissors and placed into 5 mL of 0.2 mg/mL of collagenase D in HBSS/10 mM HEPES with 5% of FBS. Then the intestinal pieces were digested at 250 rpm and 37 °C for 20 min, followed by 20 s of vortex. The resulting cell suspension contained the LPMCs, and was filtered with a 40-µm nylon cell strainer; the strainer was washed with 10 mL of PBS. LPMCs cell suspensions were spun at 500g for 10 min at 4 °C. Cell pellets were resuspended in 8 mL 44% Percoll and overlaid on 5 mL of 67% Percoll. Gradients were centrifuged at 500*g* for 20 min at 4 °C (without brake) and cells at the interface were collected and washed with 10 mL of PBS. Cells were then spun at 500*g* for 10 min at 4 °C and cells were stained for flow cytometry analysis.

### Flow cytometry and fluorescence activated cell sorting

For surface marker analysis, 1 × 10^6^ cells approximately were resuspended in 50 µL of FACS buffer. Cells were blocked with anti-CD16/CD32 for 10 min at 4 °C and then stained with the appropriate antibodies for 15 minutes at 4 °C. Antibodies and concentrations used are listed in Supplementary table 4. Cells were washed with FACS buffer, resuspended in 200 µL of FACS buffer and acquired NovoSampler Q (NovoCyte Quanteon), and absolute cell numbers and frequencies were recorded. Samples were analyzed with FlowJo (Vers.10, Tree Star) by an investigator blinded to the treatment groups (Suppl Figs 3, 4, and 9). Appropriate isotype controls, “fluorescence minus one” staining, and staining of negative populations were used to establish sorting parameters. Endothelial cells were identified as CD45^-^Ly6C^+^, microglia were identified as CD45^int^CD11b^+33^. Brain macrophages were identified as CD45^hi^CD11b^+^CD36^+26, 41, 91–93^. We could not use the canonical BAM marker CD206 because this approach requires cell permeabilization for intracellular antibody access and cannot be used in conjunction with live cells assays, such as those for measuring ROS production. Therefore, we used CD36 as a marker of BAM, as shown by single-cell RNA sequencing data^37^. To validate this approach, we quantified the overlap of CD206 and CD36 expression within the CD45 and CD11b population. We found that close to 90% of CD206 cells express CD36 (Suppl Fig 8) attesting to the validity of this approach. For fluorescence activated cell sorting, endothelial and microglia cells (Suppl Fig 7) were sorted on a FACSAria II (BD Biosciences) or CytoFlex SRT (Beckman) and collected in sample buffer for genomic DNA qRT-PCR.

### Cerebral endothelial knockdown of IL17-RA

Seven-ten week-old C57BL/6J, B6.Cg-*Gt(ROSA)26Sor^tm^*^14^*^(CAG-tdTomato)Hze^*/J (Ai14 TdTomato reporter, Stock #007909), and IL-17RA^flox/flox^ mice^34^ mice were administered 1.8×10^11^ VG in 100μL sterile PBS of AAV-BR1-iCre^32, 33^ (AAV-NRGTEWD-CAG-iCre). In Ai14 TdTomato mice, virus transduction was assessed by immunofluorescence (described above) and analyzed as TdTomato colocalization with the endothelial CD31 marker (100 vessels per mouse; n=3 mice). In IL-17RA^flox/flox^ mice, DOCA pellets were implanted three weeks after AAV-BR1-iCre administration.

### Nitric oxide measurement in pial microvessels

Pial microvessels were removed under a dissecting microscope^94^ and incubated with DAF-FM (25 μM; Molecular Probes) in l-ACSF (124 mM NaCl, 26 mM NaHCO3, 5 mM KCl, 1 mM NaH2PO4, 2 mM CaCl2, 2 mM MgSO4, 20 mM glucose, 4.5 mM lactic acid, oxygenated with 95% O2 and 5% CO2, pH=7.4) at room temperature for 45 min^24, 94^. Time-resolved fluorescence was measured every 60 s with an exposure time of 150 ms using image analysis software (IPLab, Scanalytics Inc). After a stable fluorescence baseline was achieved, microvessels were superfused with ACh (100 μM) for 15 min. DAF-FM fluorescence intensity is expressed as RFU/μm^2^, where RFU is the relative fluorescence unit, and μm^2^ is unit of the area in which RFU was measured.

### Western blotting

Cerebral blood vessels and brain microvascular endothelial cells samples were lysed in RIPA buffer (50 mM Tris-HCl pH 8.0, 150 mM NaCl, 0.5% deoxycholic acid, 0.1% SDS, 1 mM EDTA pH 8.0, 1% IGEPAL CA-630) and equal volumes were mixed with SDS sample buffer, boiled, and analyzed on 4–12% SDS polyacrylamide gels. Proteins were transferred to PVDF membranes (Millipore), blocked with 5% milk in TBS/0.1% Tween-20 (TBST) and incubated with anti-phospho-eNOS (Thr^495^) and anti-eNOS (Suppl Table 3; 1:1,000, Cell Signaling cat. #9574 and 9572, respectively). Membranes were washed in TBST, incubated with goat anti-rabbit secondary antibodies conjugated to horseradish peroxidase (1:5,000; Jackson ImmunoResearch peroxidase affinity-purified whole IgG antibodies), and protein bands were visualized with Clarity Western ECL Substrate (Bio-Rad) on a Bio-Rad ChemiDoc MP Imaging System. Raw images are provided in Suppl Fig 17-22.

### In vivo treatments

To deplete cerebral CD206^+^ brain macrophages, clodronate- or PBS-loaded liposomes were prepared as previously described^23, 33^. Under isoflurane anesthesia, 10 μl of clodronate-liposomes (7 mg/ml) and PBS-liposomes (vehicle) were injected (rate: 500 nl/min, approaching CSF clearance rates 300-400 nl/min^95^) into the cerebral ventricles (i.c.v.) with a Hamilton syringe through a burr hole drilled on the right parietal bone (coordinates: 0.5 mm posterior to bregma 1.0 mm lateral from midline, 2.3 mm below the brain surface) on the same day as DOCA pellet implantation. Assessments were made 21 days after the injection, allowing ample time for any potential perturbations in CSF homeostasis to subside. Attesting to the lack of untoward effects of the injection we did not observe any alterations in cerebrovascular regulation (functional hyperemia, endothelial vasoactivity, smooth muscle vasomotor function), BBB permeability, or cognition in mice injected with the same volume of control PBS-liposomes. FTY720 (1 mg/kg; Cayman Chemical)^24^ was injected i.p. three times every 3 d after the first week of DOCA-salt. Control and DOCA-salt mice were equipped with an intracerebroventricular (i.c.v.) cannula attached to a osmotic mini-pump (ALZET brain infusion kit 3 #0008851, pump #1004) for delivery of anti-TCRψ8 antibody (UC7-13D5, BioXCell 0.1μg/hr) or isotype control (polyclonal Armenian hamster IgG, BioXCell) on day 7 of DOCA salt. i.c.v. delivery of losartan (5μg/hour) or saline was started on the same day as DOCA pellet implantation.

### Isolation of brain and dural leukocytes

Isolation of brain leukocytes was performed as described^33^. Mice were anesthetized with pentobarbital (100 mg/kg, i.p.) and transcardially perfused with heparinized PBS. Brain cell isolation was performed by enzymatic digestion with Liberase DH (Roche Diagnostics) and Dispase (Worthington). Brain hemispheres were separated from the cerebellum and olfactory bulb and gently triturated in HEPES-HBSS buffer containing the following: 138mM NaCl, 5mM KCl, 0.4mM Na2HPO4, 0.4mM KH2PO4, 5mM d-glucose, and 10mM HEPES using a Gentle MACS dissociator (Miltenyi Biotec) following the manufacturer’s instructions. The dura was stripped from the skull^90^ using a dissection microscope. The suspension was digested with 125 μg/ml Liberase, 0.8U/ml dispase, and 50 U/ml DNase I at 37°C for 45 min (brain) or 15 min (dura) in an orbital shaker at 100 rpm. Brain cells isolated were washed and subjected to 30% Percoll (GE Healthcare) density gradient centrifugation at 500*g* for 15 min. Dura were placed on the surface of a premoistened 70-μm cell strainer. Tissue was gently homogenized with the end of a 1-mL syringe plunger, washed with 20 mL 2% FBS in PBS and centrifuged at 500*g* for 7 min.

### ROS assessment by flow cytometry

Following isolation of brain leukocytes, cells were incubated with dihydroethidium (DHE, 2.5μM) or dichlorodihydrofluorescein diacetate (DCF, 2μM) in stimulation buffer (RPMI-1640, 10% (v/v) heat inactivated FBS, 100 units/mL penicillin, 100 μg/mL streptomycin) for 30 minutes at 37° and 5% CO2. Some cells were pooled and separated for stimulation experiments, and were incubated with PBS, murine recombinant IL-17 (10ng/mL, Preprotech), or Ang II (300nM, Sigma) for 30 minutes prior to addition of DHE (as above). Cells were washed with FACS buffer (1X PBS, 2% FBS, 0.05% NaN3) and centrifuged at 500*g* for 7 min.

### Bone marrow transplant

As previously described^23, 33^, whole-body irradiation was performed on 6-week-old C57BL/6 male mice with a lethal dose of 9.5 Gy of γ radiation using a ^137^Cs source (Nordion Gammacell 40 Exactor). Eighteen hours later, BM cells (2 × 10^6^, i.v.) isolated from donor IL-17RA knockout mice or B6.129S-*Cybb^tm1Din^*/J mice (*Nox2^−/−^*, JAX stock# 002365) were transplanted in irradiated mice. Mice with transplanted BM cells were housed in cages with Sulfatrim diet for the first 2 weeks.

### qRT-PCR

Procedures for RT-PCR were identical to those previously described^23, 24, 33^. Briefly, samples were collected in TRIzol (Invitrogen Life Technologies) and RNA was extracted according to the manufacturer’s instructions. RNA samples were treated with Rnase-free DnaseI (Roche) to remove DNA contamination. cDNA was produced from mRNA samples using the iScript Reverse Transcription Supermix (Bio-Rad). Quantitative determination of gene expression was performed on a Quantstudio 5 (Applied Biosystems). Hypoxanthine-guanine phosphoribosyltransferase (HPRT) was used to normalize gene expression. qRT-PCR was conducted with cDNA in duplicate 10-μL reactions on a 384-well plate using the Maxim a SYBR Green/ROX qPCR Master Mix (2×) (Thermo Scientific). The reactions were incubated at 50°C for 2 min and then at 95°C for 10 min. A polymerase chain reaction cycling protocol consisting of 15 s at 95°C and 1 min at 60°C for 45 cycles was used for quantification. Relative expression levels were calculated according to Livak and Schmittgen, and values were normalized to respective normal control samples. Reference primer sequences are described in Suppl table 4.

For assessment of *Il17ra* knockdown *in vivo*, genomic DNA was extracted from sorted endothelial and microglia cells. Cre recombination specifically targets exons 4-7 of the *Il17ra* gene in IL17-RA^flox/flox^ mice, thus we normalized *Il17ra* exon 5 expression to the non-targeted *Il17ra* exon 3. Primer sequences as described in Suppl table 4.

### RNAscope

Tissue was treated according to manufacture’s instructions for the RNAScope Multiplex Fluorescent v2 Assay. Following 2% PFA perfusion, brain tissue was left in 2% PFA overnight. The tissue then went through a sequential sucrose gradient (10%, 20%, 30%) before freezing and storing at −80C. Tissue was cut at 10um on a cryostat, mounted on charged slides, and stored at −80C until use. Slides were left to warm up for 2 hours at −20C then baked at 60C for 30+ min. Slides were washed with 1x PBS for 5 min then treated with RNAScope Hydrogen Peroxide for 10 min. Slides were washed with nuclease-free water and pretreated with target retrieval buffer in a slide envelope placed in 100C boiling water for 5 min. Slides were then washed with distilled water, dipped in 100% EtOH, and a barrier was established around the tissue using a hydrophobic marker. Slides were then allowed to air dry overnight. Slides were treated with Protease IV for 15 min at 40C, followed by a probe incubation for 2 hours at 40C. The probes (purchased from ACD) used were Mm-Mrc1 (Cat # 437511) and Mm-Il17ra-O1-C2 (Cat # 566131-C2). A 3-plex positive control probe and 3-plex negative control were run concurrently. After probe incubation, slides underwent three amplification steps (RNAScope Multiplex Fluorescent Detection Reagents v2, ACD, Cat # 323110) followed by the development of horseradish peroxidase (HRP) signal fluorescence with TSA-based fluorophores. HRP blockers were used after every fluorophore incubation. After each step, slides were washed twice with wash buffer. Slides were then treated with RNAScope DAPI for 30 seconds then cover-slipped in Fluorosave Reagent (Sigma, Cat # 345789) and left to dry for 20 min. Slides were then stored at 4C until imaging.

### IL-17A ELISpot Assay

Dura leukocytes were isolated as described above, and incubated in ELISpot plate for 4 hours in stimulation buffer (RPMI-1640, 10% (v/v) heat inactivated FBS, 100 units/mL penicillin, 100 μg/mL streptomycin, 100 ng/mL phorbol 12-myristate 13-acetate (PMA), 1 µg/mL of ionomycin) in a 37C humidified incubator with 5% CO2. Mouse IL-17A ELISpotPLUS kit ALP (MABTECH, Cat# 3521-4APW-2) was performed following manufacturer’s instructions. Wells were developed with BCIP/NBT-plus for 15 minutes, and color development was stopped by washing with diH2O. Images were obtained using a digital camera (T TAKMLY, MX200-B) and analyzed using ImageJ software.

### CSF Collection

Mice were anesthetized with isoflurane (3% induction, 1.5% maintenance), and placed on a stereotaxic frame with the head slightly tilted forward. A small incision was made over the back of the neck, and muscle tissue moved to expose the dura over the cisterna magna. The area was cleaned with a cotton bud. Heparinized glass capillaries were prepared using a micropipette puller, attached to a thin tube and syringe connected through a three-way valve, and mounted on a micromanipulator^96^. Under a dissecting microscope, the glass capillary was aligned to the cisterna magna, avoiding blood vessels in the area, and tapped through the dura to collect CSF for a maximum of 10 minutes. On average, 15-18μl was obtained per mouse. Equal volumes of CSF from two animals were pooled to maximize loading sample volume of 25μl for IL-17 quantification (Mesoscale).

### Ang II assay

Mice were euthanized, and blood was collected by cardiac puncture in a prechilled tube containing a mix of protease inhibitors. Plasma was collected and transferred to a prechilled tube and stored frozen at –80°C. Brains were collected and immediately frozen on dry ice. Ang II concentration was determined using a radioimmunoassay by the Hypertension Core Laboratory of the Wake Forest University School of Medicine (Winston-Salem, North Carolina, USA)^97^.

### Statistical analysis

Sample size was determined according to standard power analysis (80% power and α=0.05) based on standardized effect sizes calculated from published studies^23, 24, 30, 88, 98^. Animals were randomly assigned to treatment and control groups, and analysis was performed in a blinded fashion. After testing for normality (Bartlett’s test), intergroup differences were analyzed by unpaired two-tailed t-test for single comparison, or by 1-way ANOVA or 2-way ANOVA with Tukey’s or Bonferroni’s multiple comparison test, as appropriate and indicated in the figure legends. If non-parametric testing was indicated, intergroup differences were analyzed by Mann-Whitney test or Kruskal-Wallis test with Dunn’s correction, as appropriate and indicated in the figure legends. Statistical tests through the manuscript were performed using Prism 7 (GraphPad). All data are presented as mean ± SEM.

## NONSTANDARD ABBREVIATIONS

Ach: acetylcholine
Ang II: angiotensin II
ASL: arterial spin label
AT1R: ang II type 1 receptor
BAM: brain associated macrophages
bECKO: brain endothelial cell knockout
BM: bone marrow
BP: blood pressure
CBF: cerebral blood flow
DOCA: deoxycorticosterone acetate
eNOS: endothelial nitric oxide synthase
HTN: hypertension
i.c.v.: intracerebroventricular
IL-17: interleukin 17
IL-17RA: IL-17 receptor A
KO: knockout
NO: nitric oxide
RAS: renin angiotensin system
ROS: reactive oxygen species
Th17: T-helper 17
WT: wild-type

## ACKNOWLEDGEMENTS

This work was supported by grants R37-NS089323 (CI), R01-NS095441(CI) and K22-NS123507 (MMS), as well as the Leon Levy Fellowship in Neuroscience (MMS). The support from the Feil Family Foundation is gratefully acknowledged. The authors declare no competing financial interests. Figure schematics were made on Biorender.

## Notes

### Competing Interest Statement

The authors have declared no competing interest.

### Summary of Updates

New figure 5, new data in figures 6 and 7. Updated supplemental files.

