## Supplement for "Meningeal IL-17 producing T cells mediate cognitive impairment in salt-sensitive hypertension"

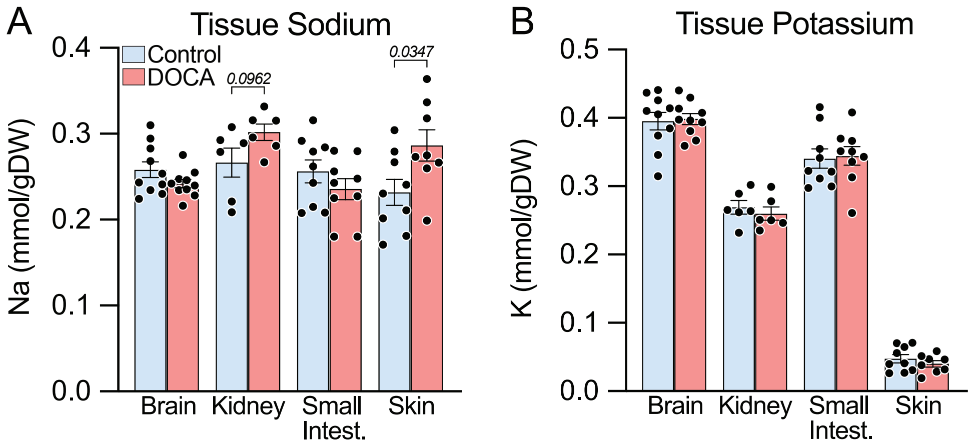


**Supplementary Figure 1.** Tissue sodium and potassium content at 21 days of DOCA-salt HTN.
**(A and B)** Tissue sodium and potassium content was assessed by inductively coupled plasma – atomic emission spectrometry (ICP-AES)[75, 76]. Intergroup differences analyzed by unpaired two-tailed t-test for each organ.

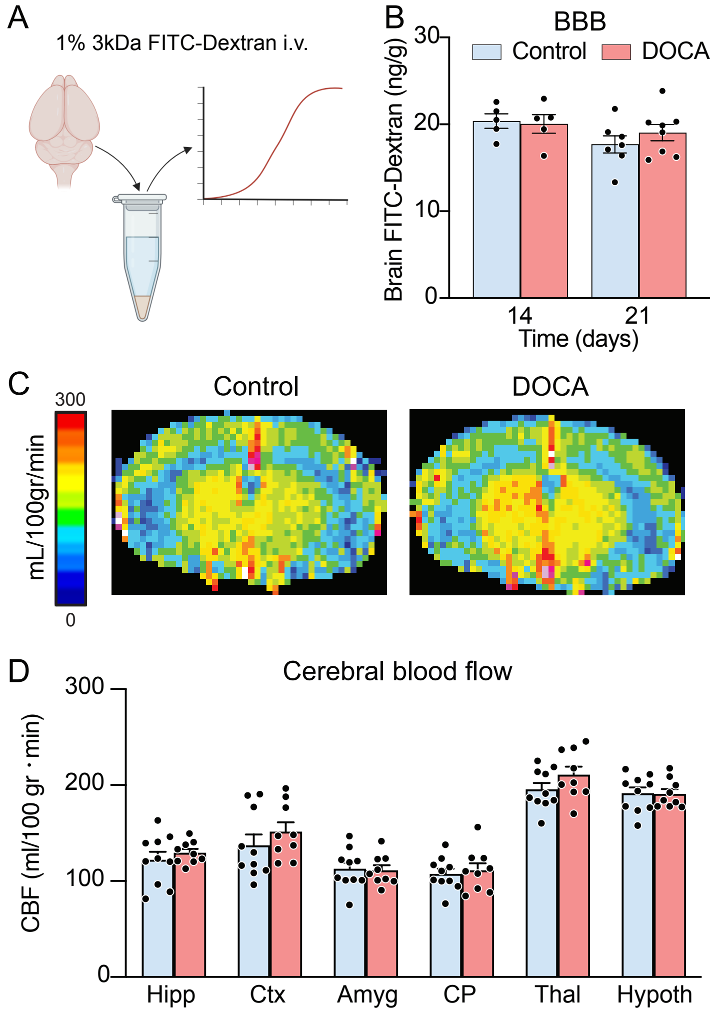


**Supplementary Figure 2.** Blood-brain barrier (BBB) integrity and resting cerebral blood flow (CBF) are not impaired by DOCA-salt. **(A and B)** BBB permeability assessed by brain extravasation of 3kDa FITC-dextran quantified by spectrophotometry in brain homogenates revealed no impairment during DOCA-salt hypertension (n=5-8). **(C and D)** DOCA-salt hypertension does not impair resting CBF assessed quantitatively by arterial spin label (ASL)-MRI (n=9-10) at 21 days of treatment in the hippocampus (Hipp), cortex (Ctx), amygdala (Amyg), caudate putamen (CP), thalamus (Thal), or hypothalamus (Hypoth). Intergroup differences analyzed by two-way ANOVA with Tukey's multiple comparisons test.

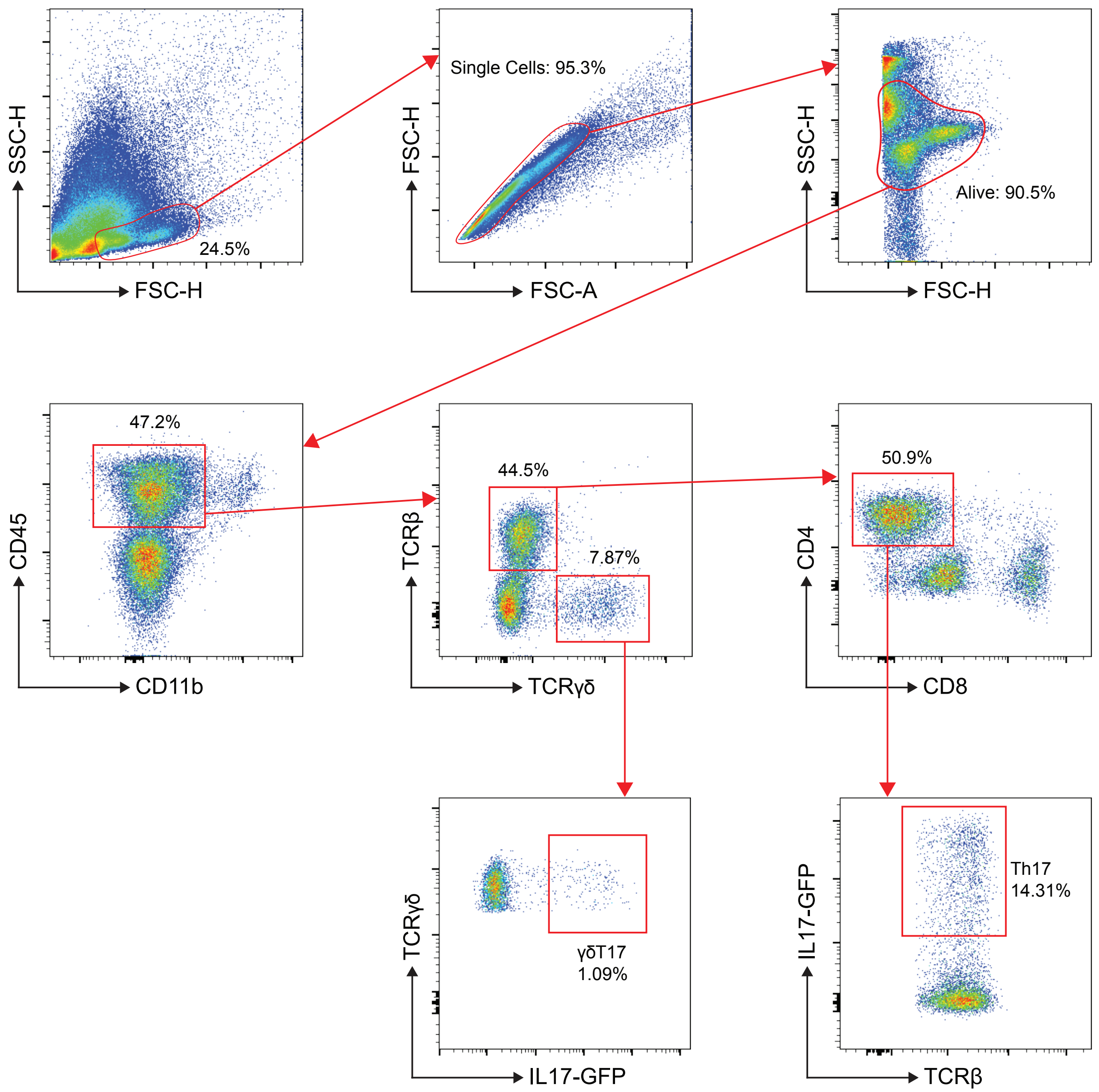


**Supplementary Figure 3.** Gating strategy for Th17 and γδT17.

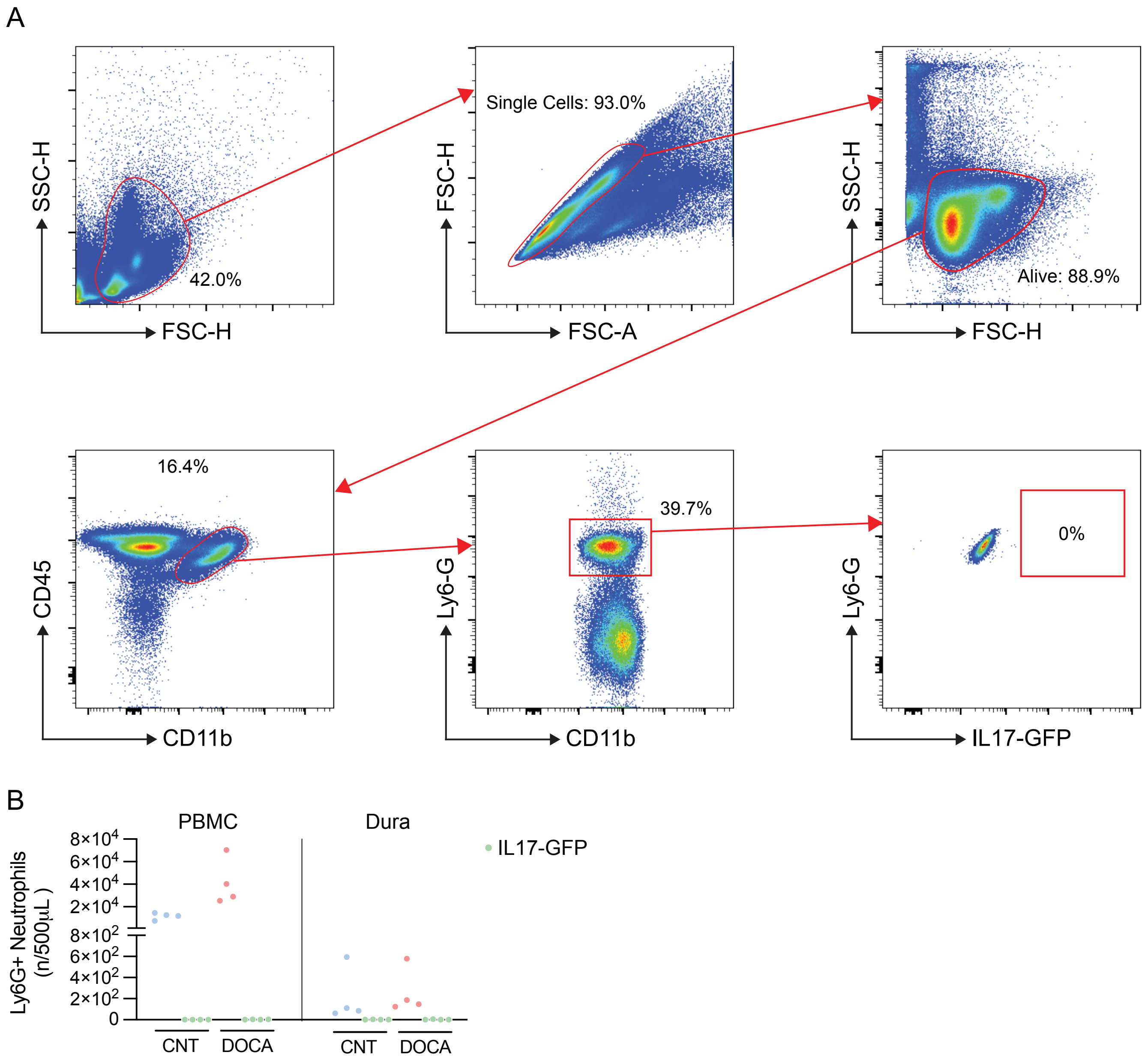


**Supplementary Figure 4.** Neutrophils do not increase IL-17 production following DOCA-salt. **(A)** Gating strategy for identification of neutrophils ($CD45^+CD11b^+Ly6\text{-}G^+$). **(B)** IL17-GFP+ neutrophils were not changed by DOCA-salt in peripheral blood mononuclear cells (PBMC) or in the dura. n=4/group.

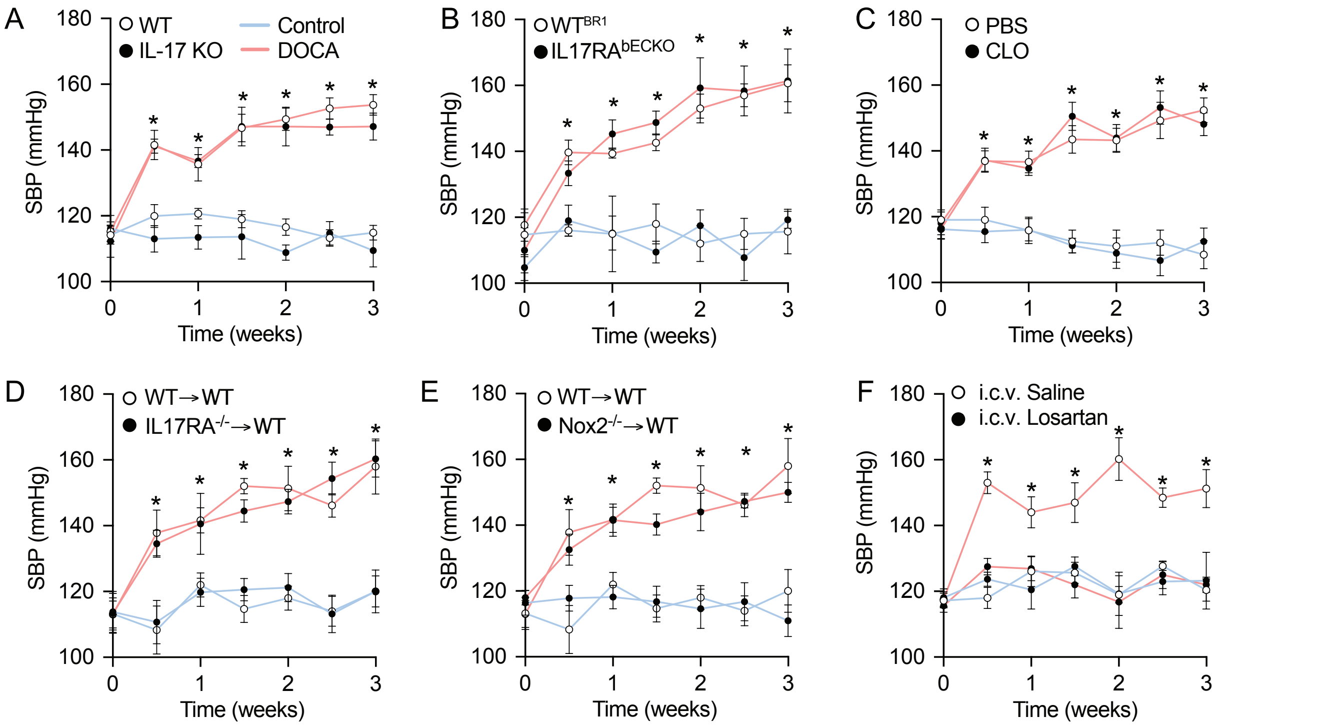


**Supplementary Figure 5.** Systolic blood pressure measured by tail-cuff plethysmography. Systolic blood pressure (SBP) was assessed twice per week in all control and DOCA-salt mice of the following groups: **(A)** wild-type (WT) and IL17-deficient (IL17KO) mice, **(B)** WT and IL17RA brain endothelial cell knockout ($IL17RA^{bECKO}$), **(C)** mice treated with vehicle (PBS) or clodronate (CLO)-containing liposomes, **(D and E)** bone marrow chimeras, and **(F)** i.c.v. saline or losartan. Intergroup differences analyzed by two-way repeated measures ANOVA with Tukey's multiple comparisons test.

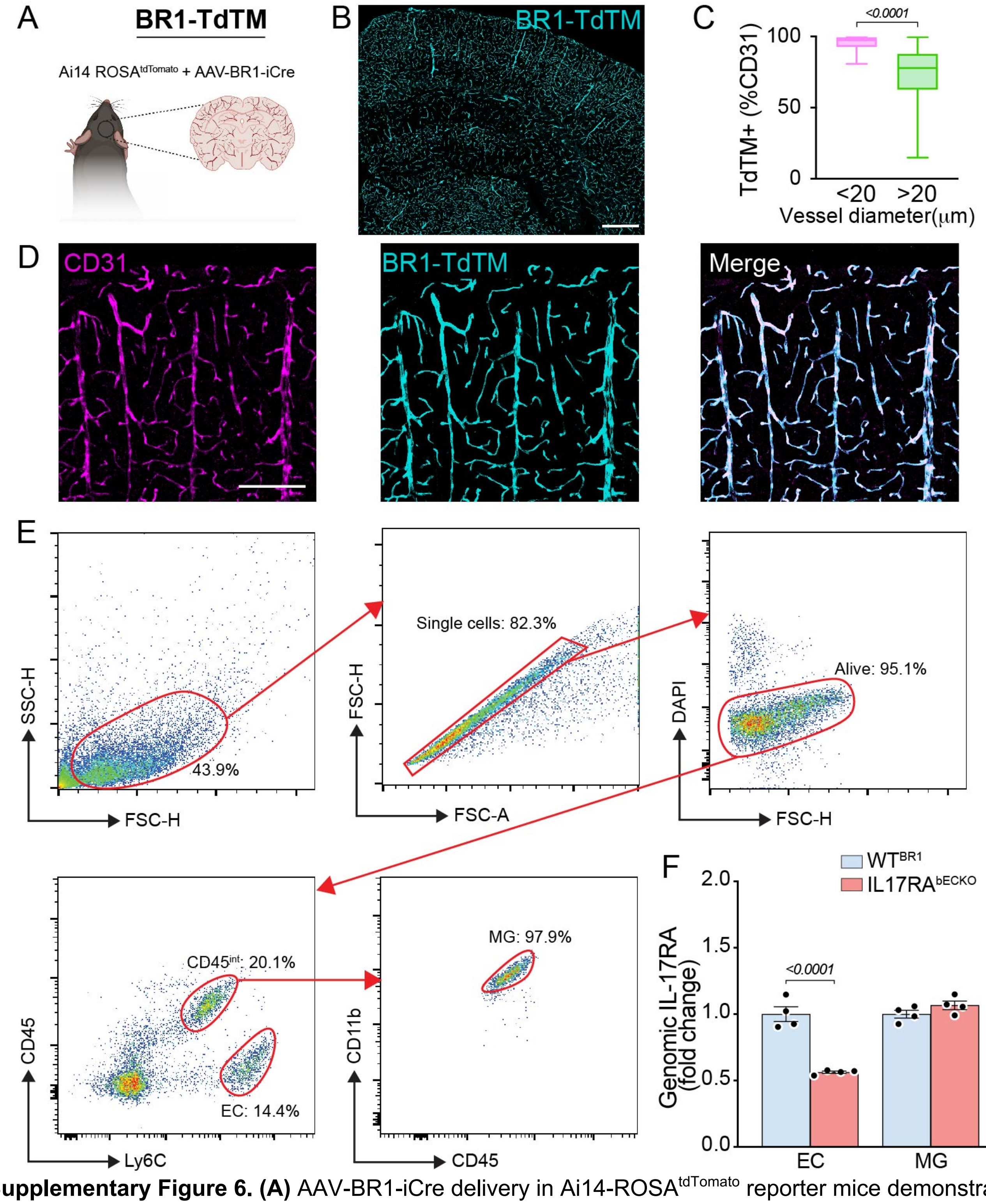


**Supplementary Figure 6. (A)** AAV-BR1-iCre delivery in Ai14-ROSA$^{tdTomato}$ reporter mice demonstrates **(B)** widespread TdTomato (TdTM) expression in cerebral vessels. Scale bar: 500μm. **(C)** Specifically, we observed a 90-95% endothelial viral transduction in vessels less than 20μm (n=3 mice, 100 vessels per mouse). **(D)** Representative images of TdTM expression in CD31+ endothelial cells. Scale bar: 150μm. **(E)** Sorting strategy for assessing IL-17RA gene deletion in IL17RA brain endothelial cell knockout (bECKO) mice. Endothelial cells were identified as CD45$^{-}$Ly6C$^{+}$, microglia were identified as CD45$^{int}$CD11b$^{+}$[32]. **(F)** Quantification of genomic IL-17RA deletion in EC and MG. Intergroup differences analyzed by two-way ANOVA with Tukey's multiple-comparison test.

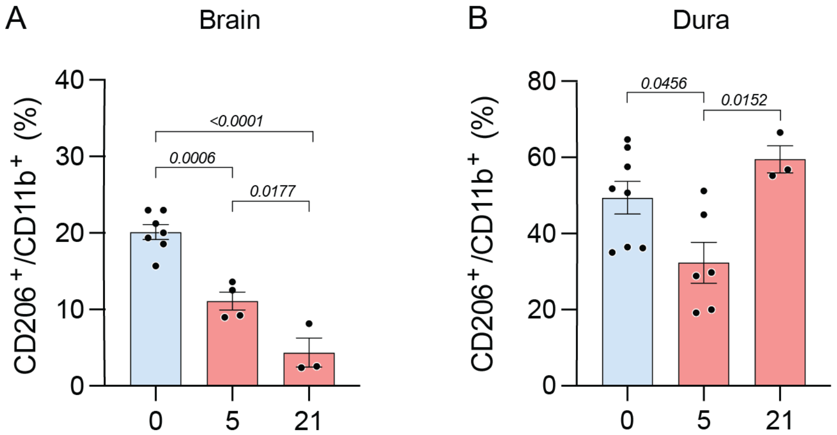


**Supplementary Figure 7. (A)** i.c.v. clodronate depletes brain macrophages for 21 days, and **(B)** initially depletes dura macrophages, but they are fully restored within 21 days. n=3-7/group. Intergroup differences analyzed by one-way ANOVA with Tukey's multiple-comparison test.

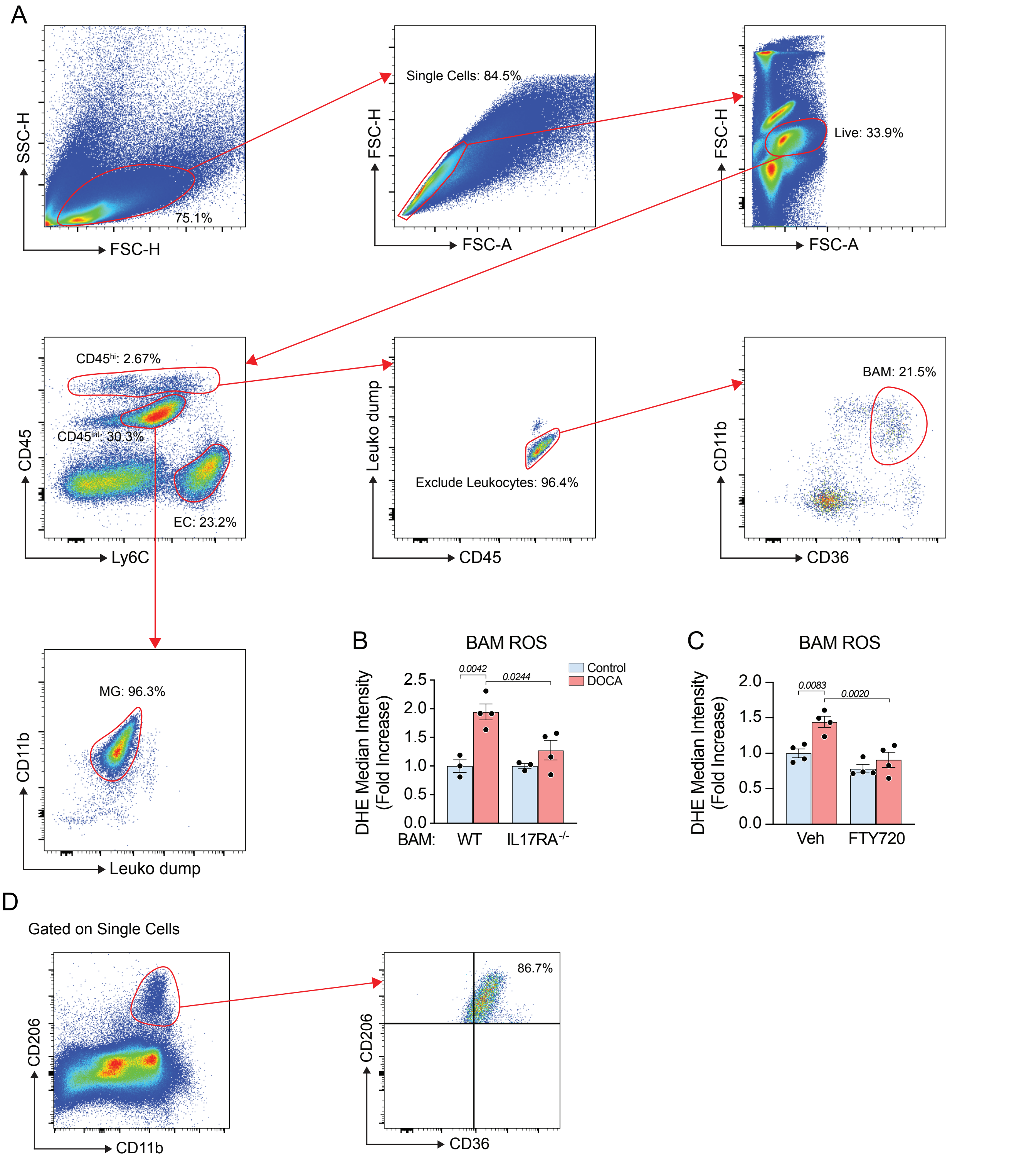


**Supplementary Figure 9. (A)** Gating strategy for identification of endothelial cells (EC), microglia (MG), and brain associated macrophages (BAM) for assessment of reactive oxygen species (ROS). For the exclusion of blood derived leukocytes (Leuko), the “Leuko dump” channel included: CD3e, CD19, NK1.1, TCRβ, and TCRγδ. **(B)** DOCA-salt does not increase BAM ROS IL17RA$^{-/-}$→WT chimeras, or **(C)** in WT mice treated with FTY720. Intergroup differences analyzed by two-way ANOVA with Tukey’s multiple comparisons test. **(D)** Validation of CD36 to identify BAM, following traditional BAM gating strategy of CD206+ and CD11b+, 86.7% of these cells are CD36+.

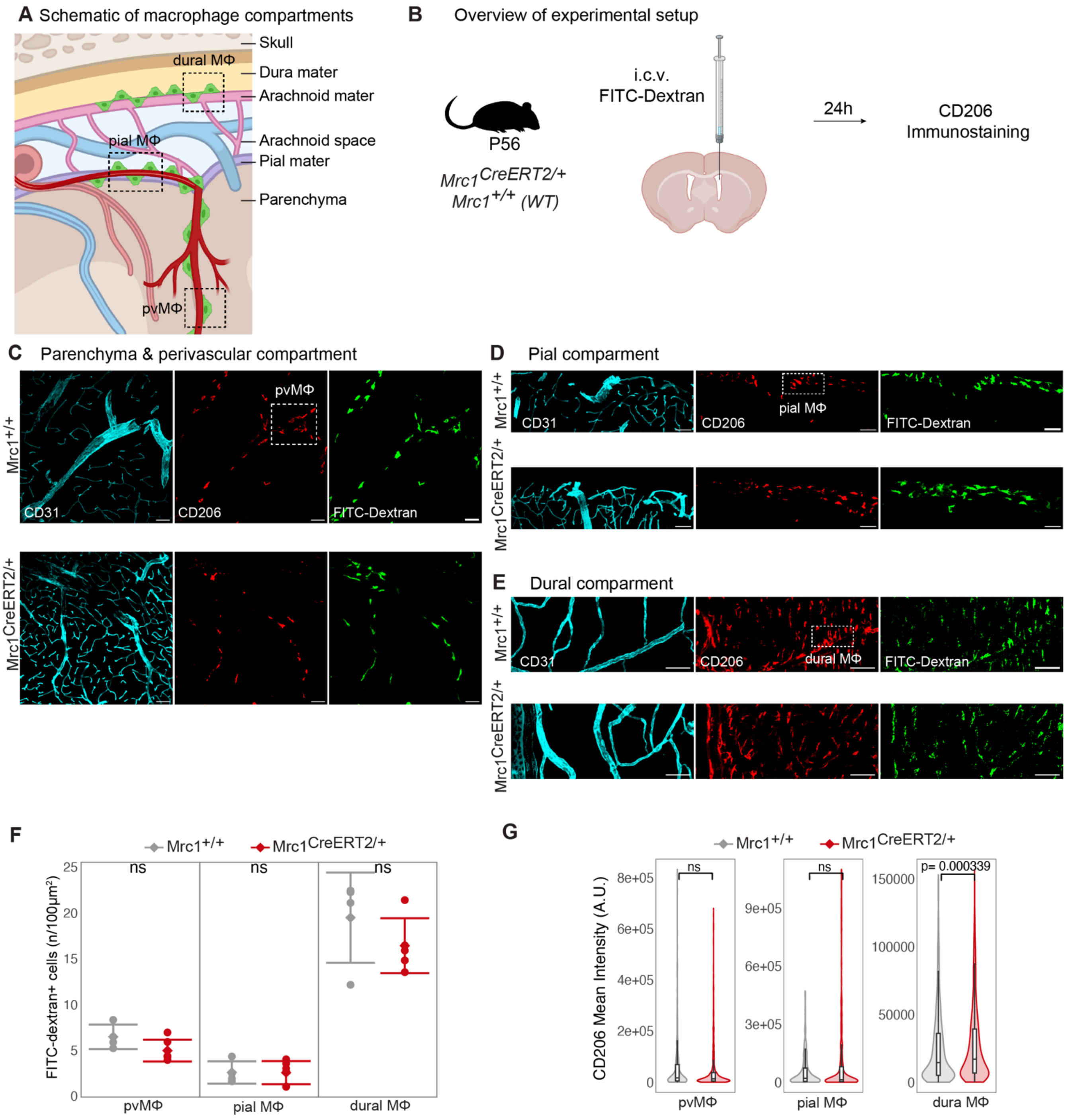


**Supplementary Figure 10. Novel Mrc1$^{CreERT2/+}$ mouse.**
**(A)** Schematic of CNS macrophage compartments. **(B)** B. Illustration of the experimental procedure for intracerebroventricular (i.c.v.) FITC-Dextran injection and analysis of adult Mrc1$^{+/+}$ or Mrc1$^{CreERT2/+}$. **(C-E)** Immunofluorescence images reveals FITC-Dextran (green) in BAMs (CD206+, red) in perivascular macrophages (pvMΦ) and pial MΦ and dural MΦ compartments in the cortex of adult Mrc1$^{+/+}$ or Mrc1$^{CreERT2/+}$ mice. CD31+ blood vessels shown in cyan. Scale bars: 20 µm. **(F)** Quantification of pvMΦ and pial MΦ and dural MΦ in *Mrc1$^{+/+}$* or *Mrc1$^{CreERT2/+}$* mice. **(G)** CD206 surface expression levels on individual FITC-Dextran cells. p-values determined by Wilcoxon signed rank test. Data shown as mean ± SEM; n=4-5/group.

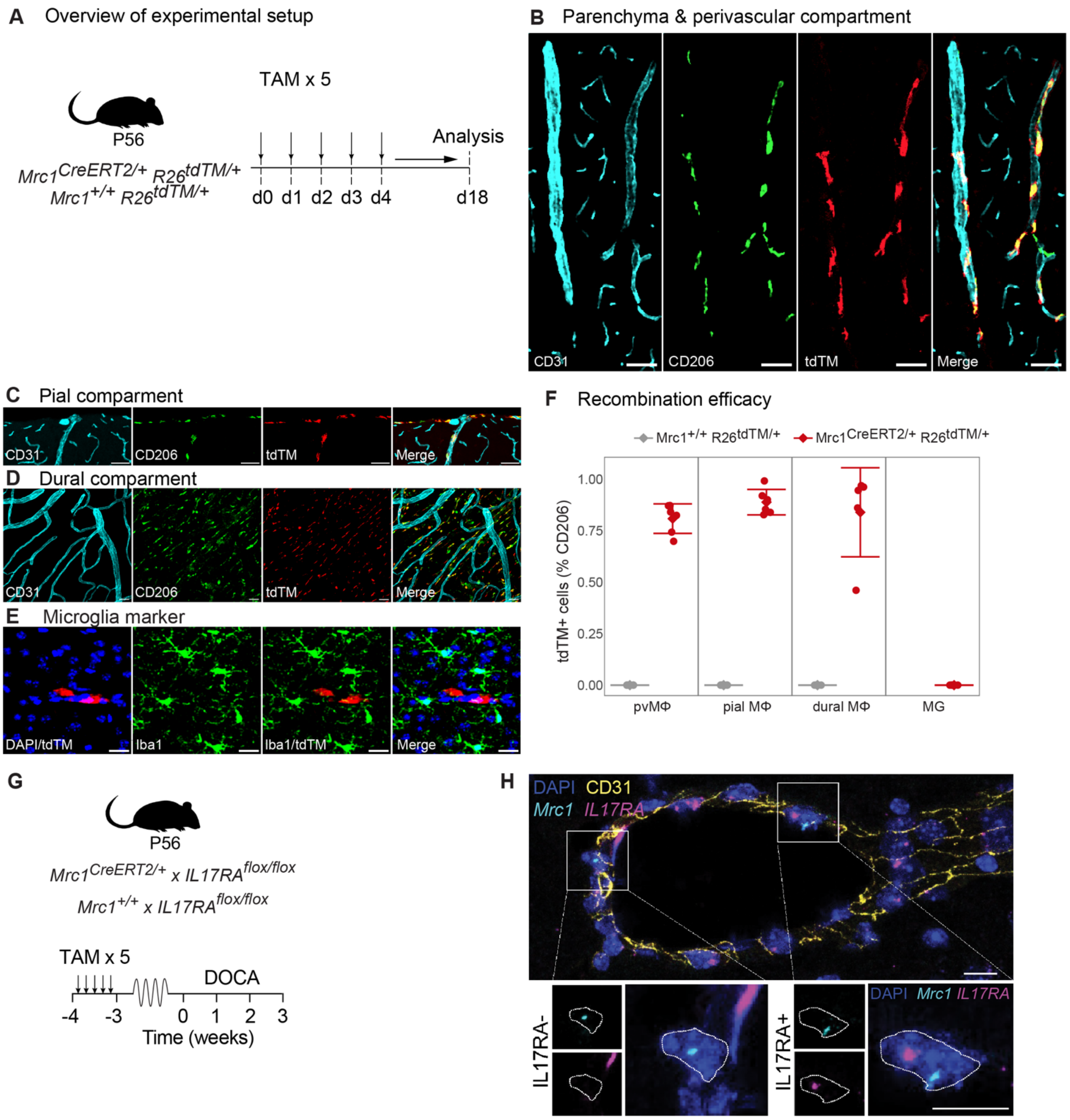


**Supplementary Figure 11. Novel Mrc1CreERT2/+ mouse.**

**(A)** Experimental procedure for tamoxifen administration (TAM) and analysis of adult *$Mrc1^{+/+}R26^{tdTM/+}$* or *$Mrc1^{CreERT2/+}R26^{tdTM/+}$* mice. **(B-D)** Immunofluorescence images reveals $tdTM^{+}$ (red) in $CD206^{+}$ BAM (green) in perivascular macrophages (pvMΦ), pial MΦ, and dural MΦ. $CD31^{+}$ blood vessels shown in cyan. Scale bars: 20 µm**. (E)** TdTM expression was no observed in microglia ($IBA1^{+}$, green) of adult *$Mrc1^{CreERT2/+}R26^{tdTM/+}$* mice. Scale bars: 20 µm**. (F)** Quantification of recombination efficacy in pvMΦ, pial MΦ, and dural MΦ, as well as microglia in *$Mrc1^{+/+}R26^{tdTM/+}$* and *$Mrc1^{CreERT2/+}R26^{tdTM/+}$* mice. Data shown as mean ± SEM; n=4-5/group. **(G)** Experimental procedure for tamoxifen administration and analysis of $Mrc1^{CreERT2/+}$ x $IL17RA^{flox/flox}$ mice. **(H)** Representative image of cerebral blood vessel stained for CD31 (yellow, IHC), Mrc1 (cyan, RNAscope) and IL17RA (magenta, RNAscope). Blood vessel shows one IL17RA- BAM and one IL17RA+ BAM. This identification strategy was used for quantification of IL17RA deletion in $Mrc1^{CreERT2/+}$ x $IL17RA^{flox/flox}$ mice. Scale bars: 10 µm**.**

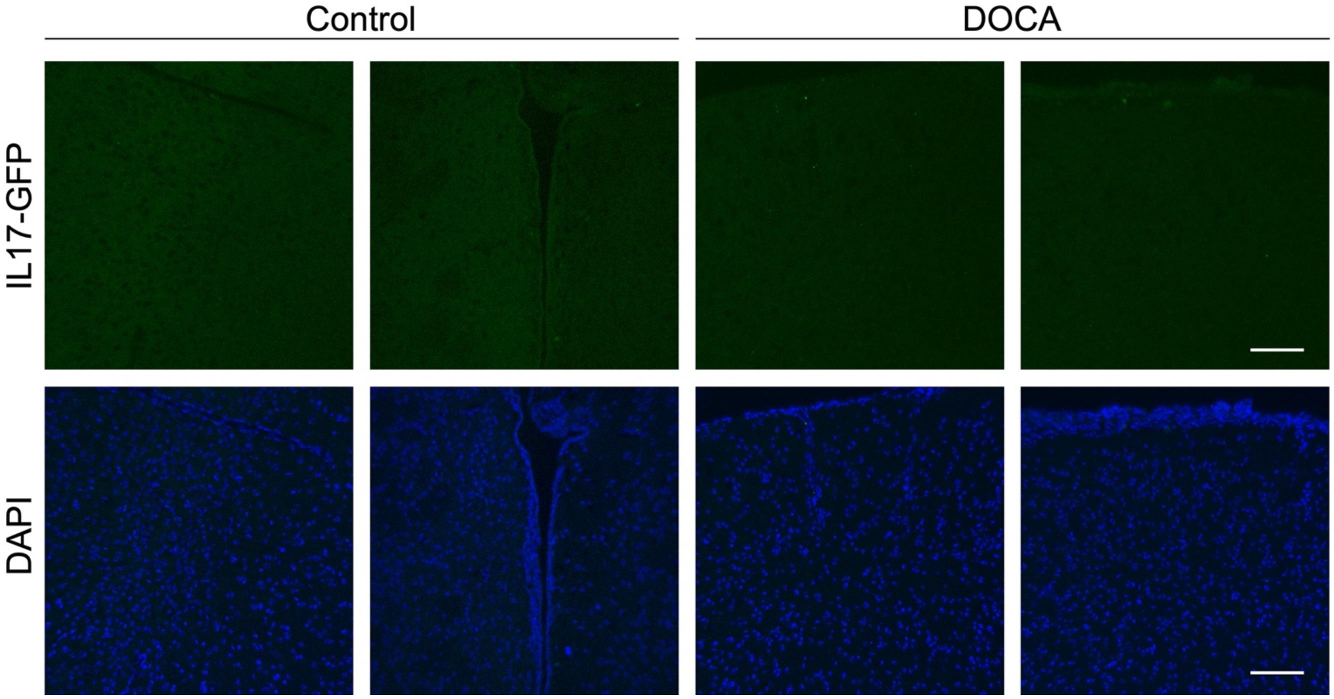


**Supplementary Figure 12.** Representative images of brain tissue from control and DOCA-salt IL17-GFP mice. IL17-GFP cells were not observed in the brain parenchyma. At least 10 sections from n=6 mice per group were examined. Scale bars: 100μm.

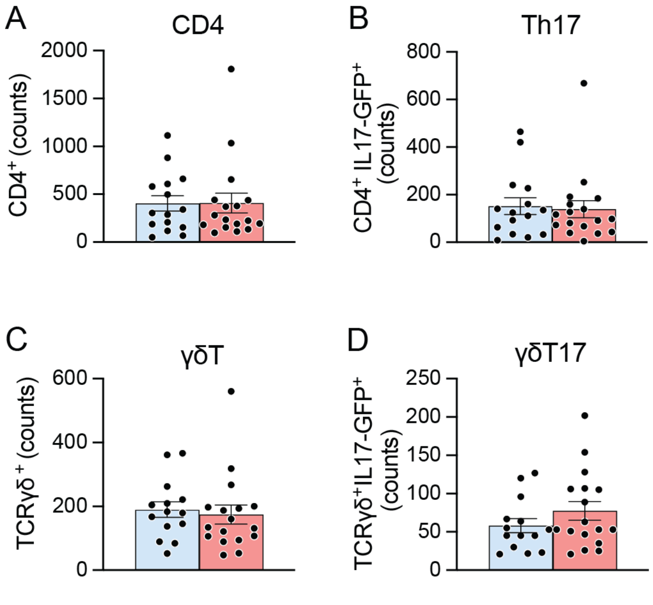


**Supplementary Figure 13.** Total numbers of cells obtained by flow cytometry from control and DOCA dura samples. **(A)** Total CD4 cells. **(B)** Total Th17 cells. **(C)** Total γδT cells. **(D)**Total γδT17 cells.

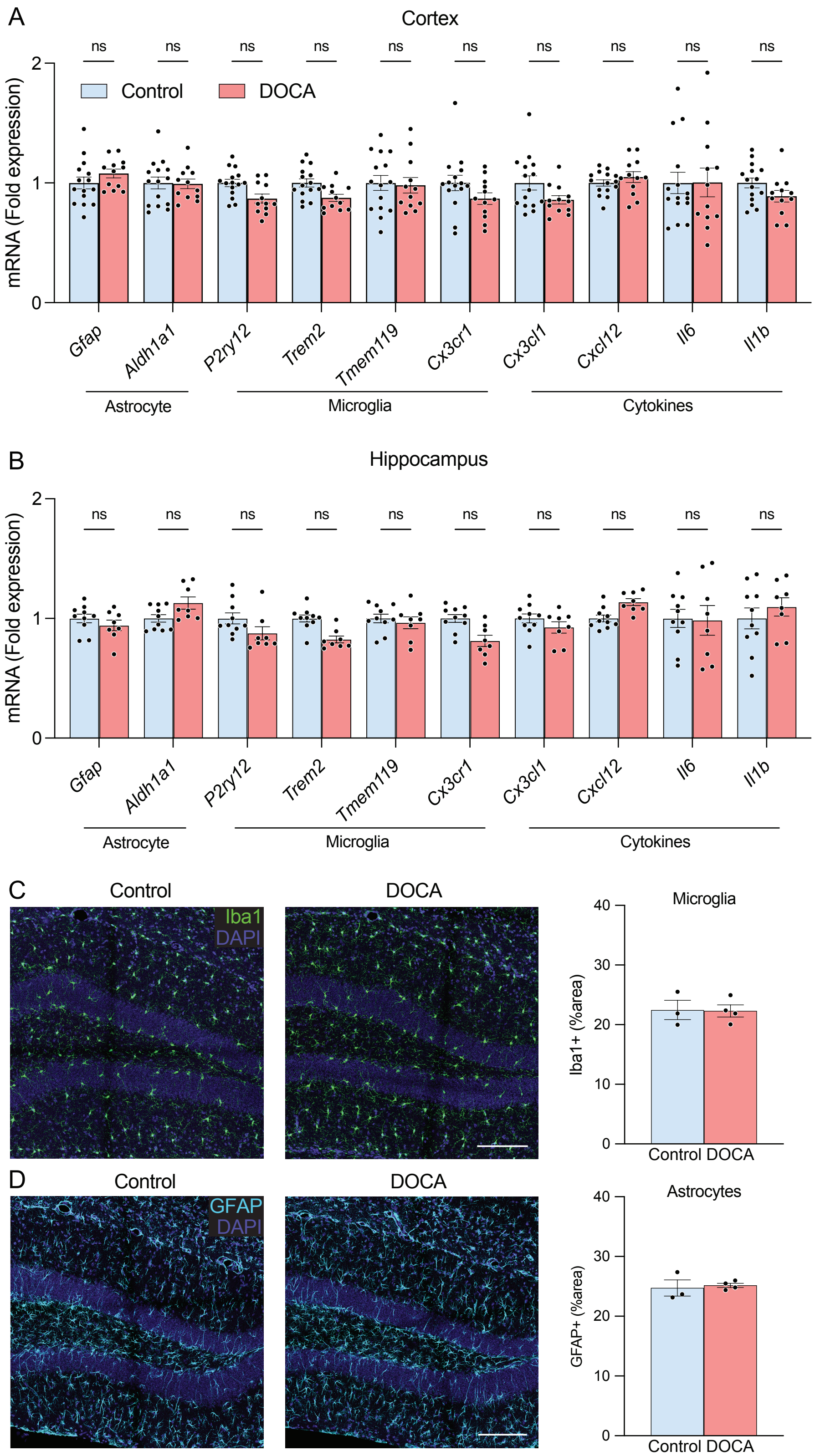


**Supplementary Figure 14.** Markers of neuroinflammation in neocortex and hippocampus. **(A-B)** Selected inflammatory gene expression assessed by qPCR was not altered in cortex or hippocampus from DOCA-salt mice. Intergroup differences analyzed by two-way ANOVA with Tukey's multiple comparison test. **(C)** Iba1+ microglia and **(D)** GFAP+ astrocyte area was not altered in the hippocampus of DOCA-salt. Intergroup differences analyzed by unpaired two-tailed t-test. Scale bars: 150μm.

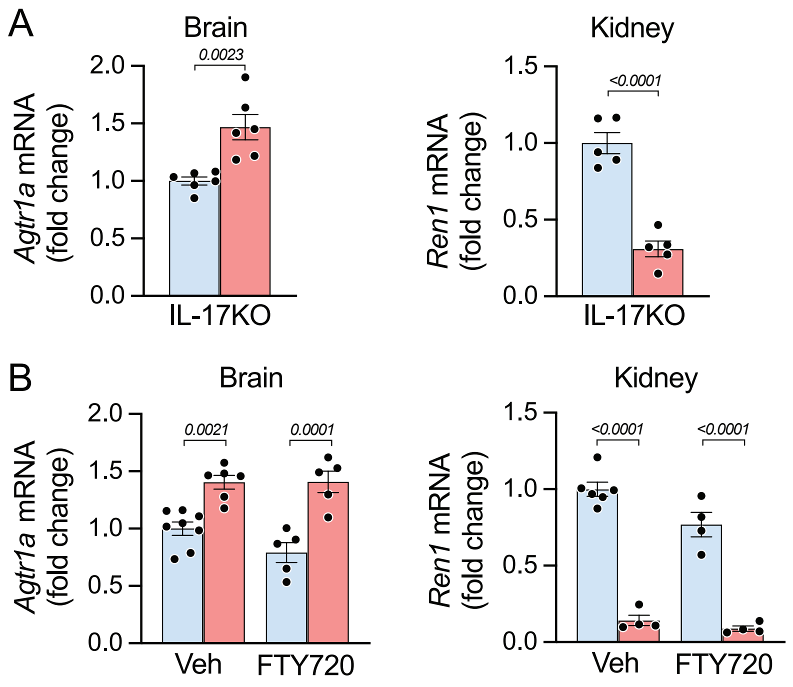


**Supplementary Figure 15.** Brain *Agtr1a* and renal *Ren1* mRNA expression in **(A)** IL-17KO and **(B)** vehicle and FTY720-treated control and DOCA-salt mice. Intergroup differences analyzed by unpaired two-tailed t-test (A) or two-way ANOVA with Tukey's multiple comparison test (B).

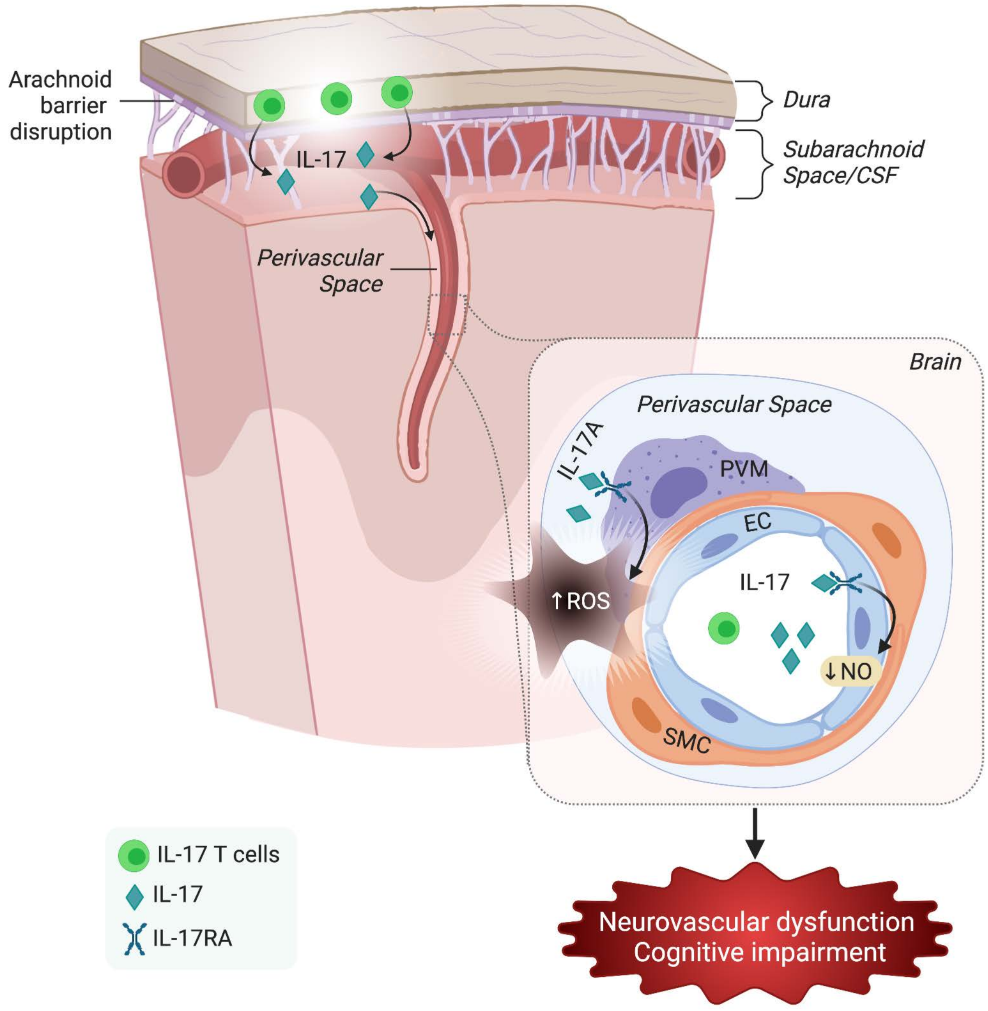


**Supplementary Figure 16.** Summary of the mechanisms by which DOCA-salt hypertension alters neurovascular and cognitive function in mice. These effects are mediated by concurring actions of IL17 acting on IL17RA on different cells types on both sides of the BBB. In the circulation, IL-17 produced by T-cells acts on cerebral endothelial IL-17RA to reduce NO production leading to suppression of endothelial vasoactivity without affecting the increase in CBF induced by neural activity. In the brain, IL-17 produced by dura T-cells acts on IL-17RA on BAM to induce vascular oxidative stress and suppression of functional hyperemia with minimal effects on endothelial function.

211227-vessels22A-t495-MJS_05

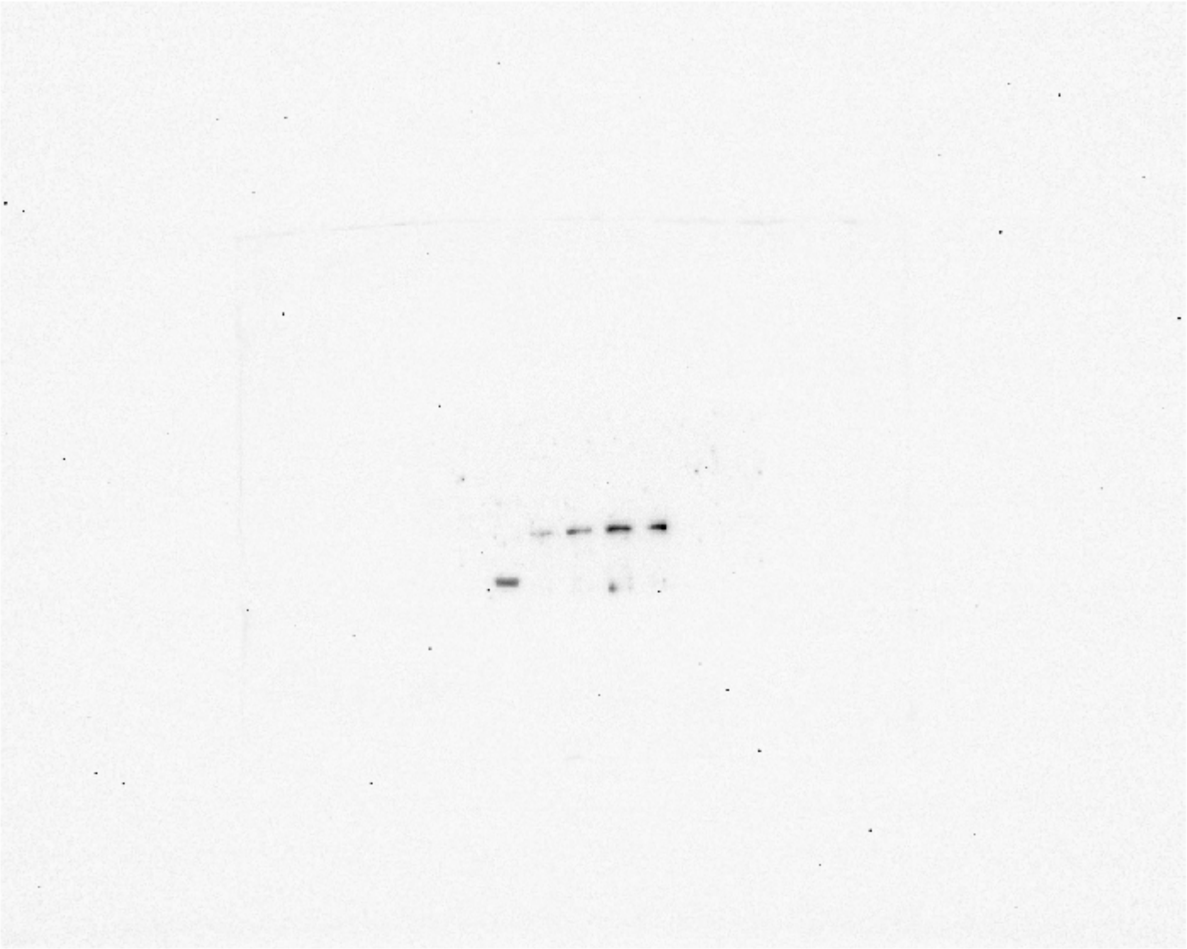

**Supplementary Figure 17.** Raw western blot image pThr495 in Fig. 3E.

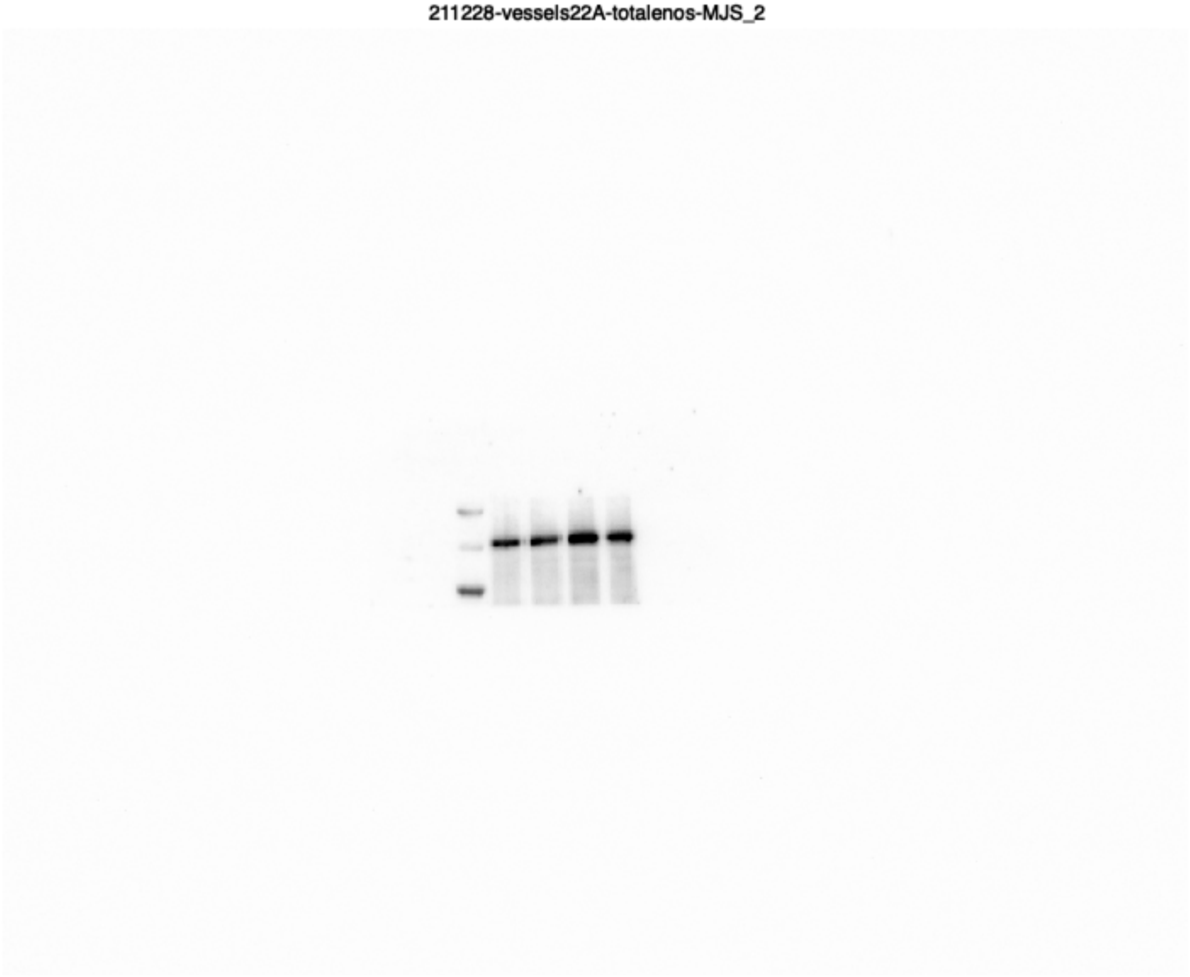


**Supplementary Figure 18.** Raw western blot image total eNOS in Fig. 3E.

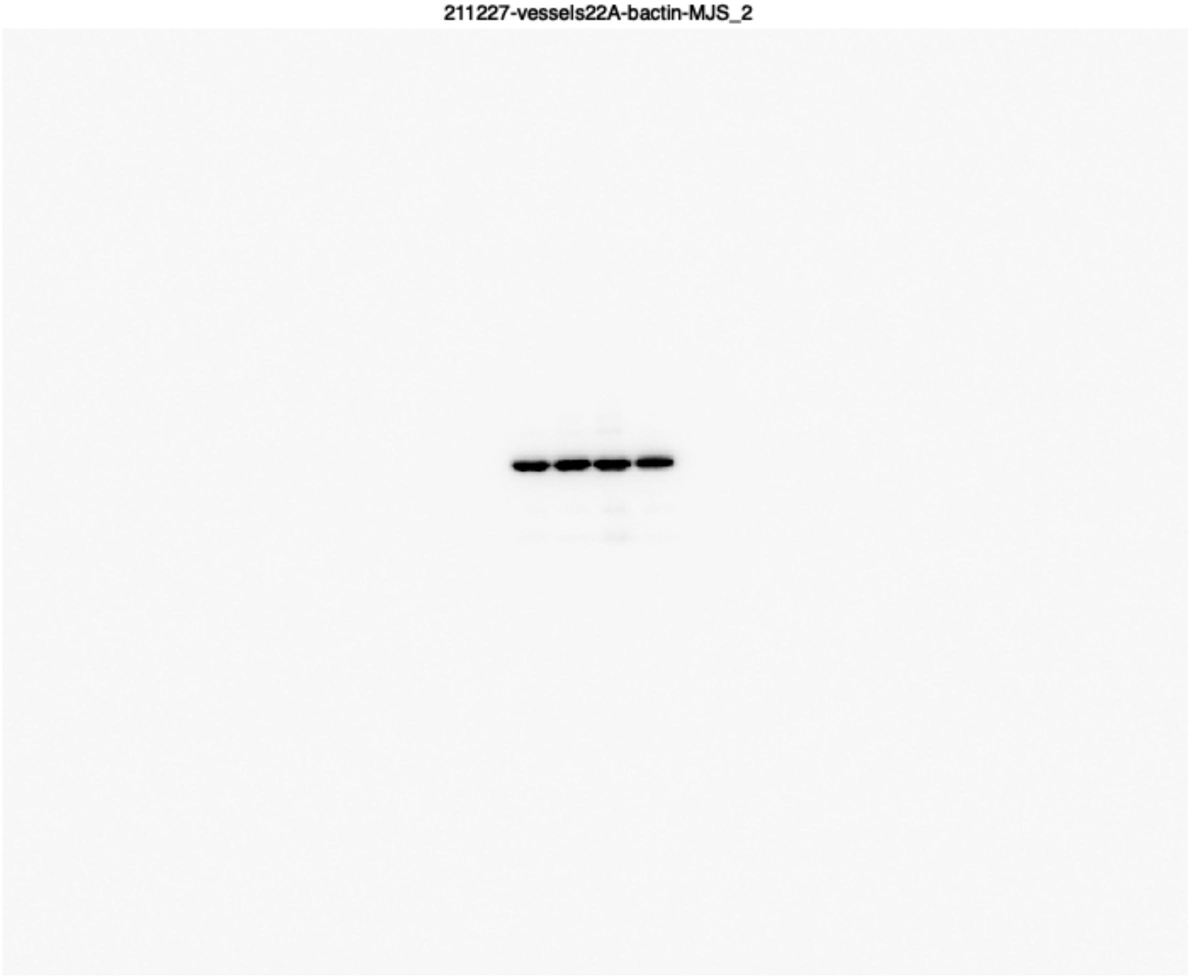


**Supplementary Figure 19.** Raw western blot image beta-actin in Fig. 3E.

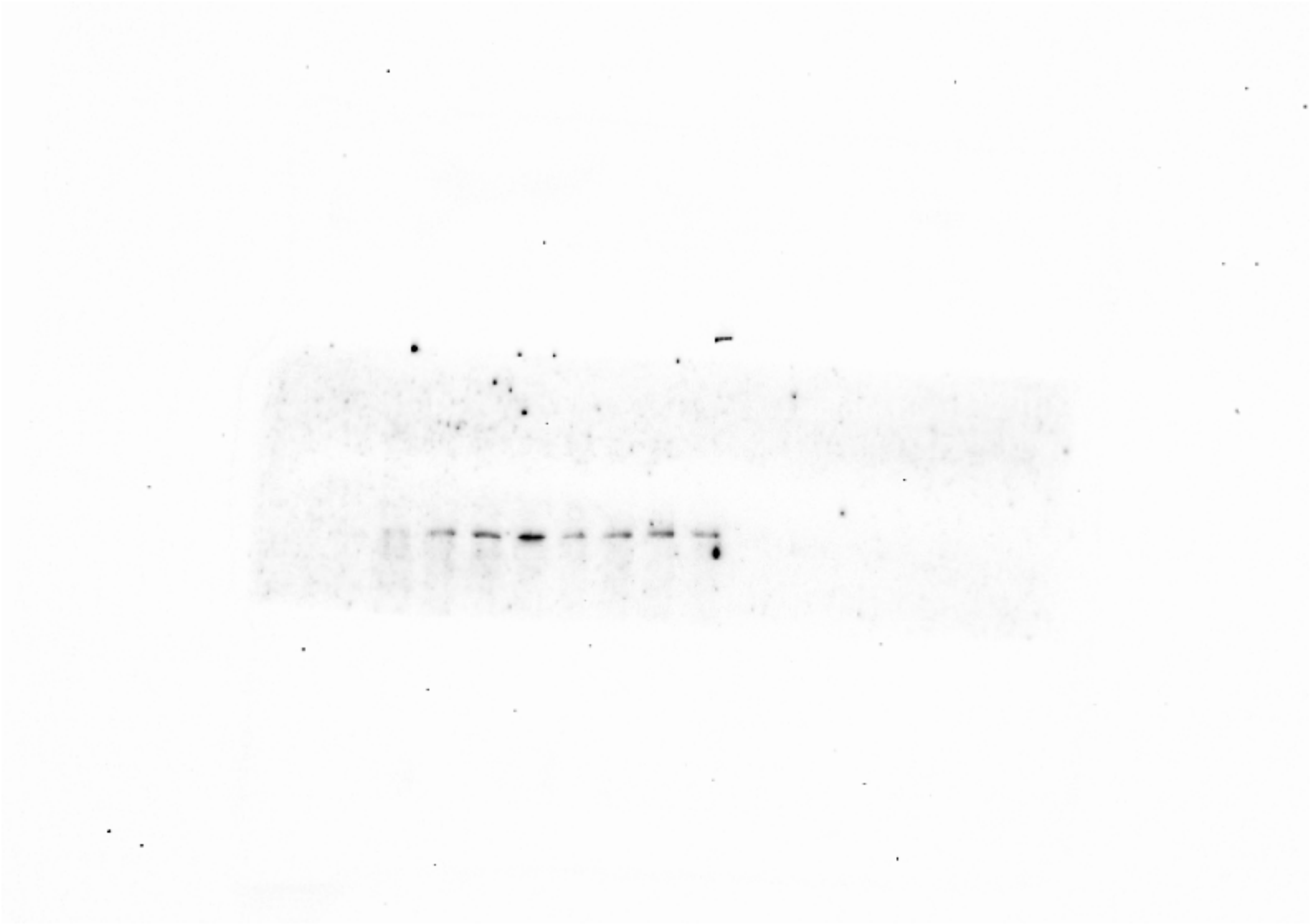

**Supplementary Figure 20.** Raw western blot image pThr495 in Fig. 3F.

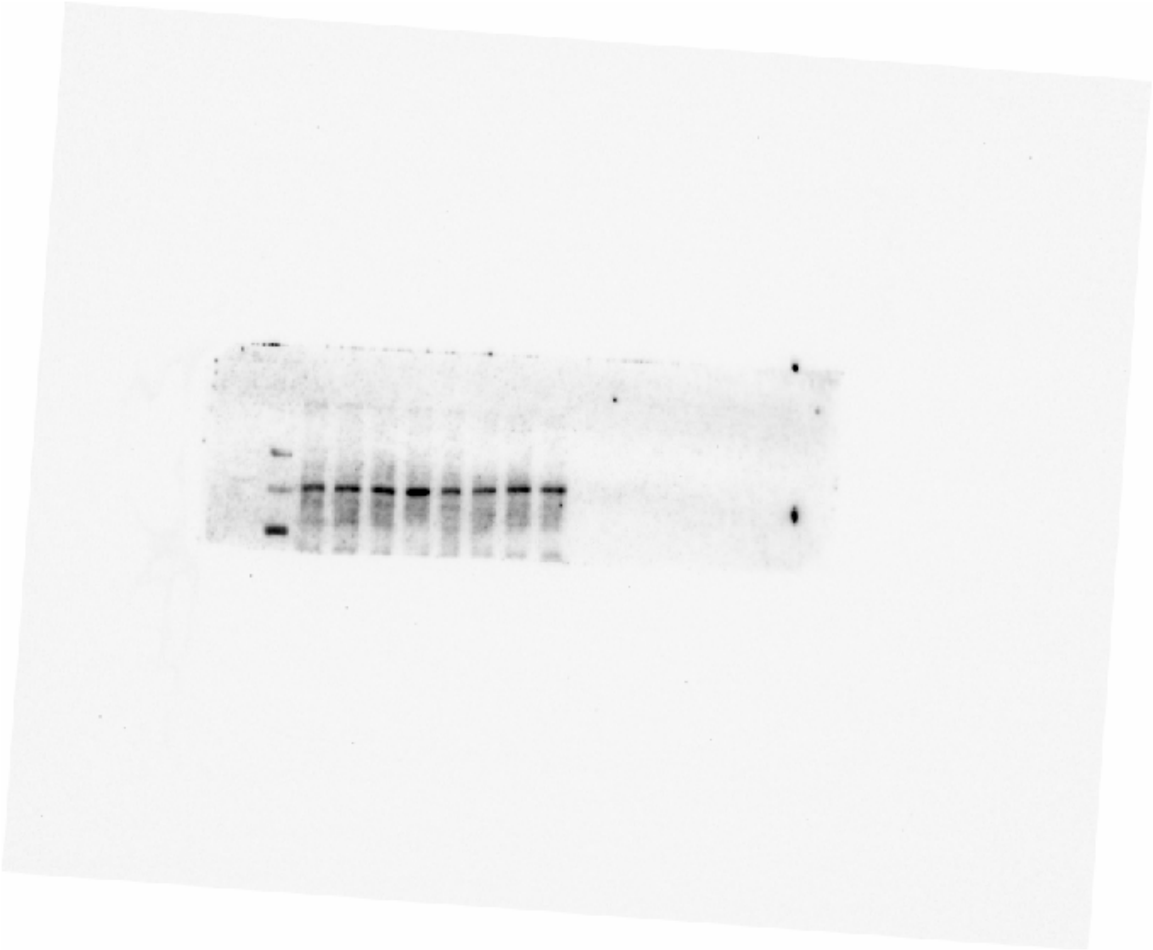

**Supplementary Figure 21.** Raw western blot image total eNOS in Fig. 3F.

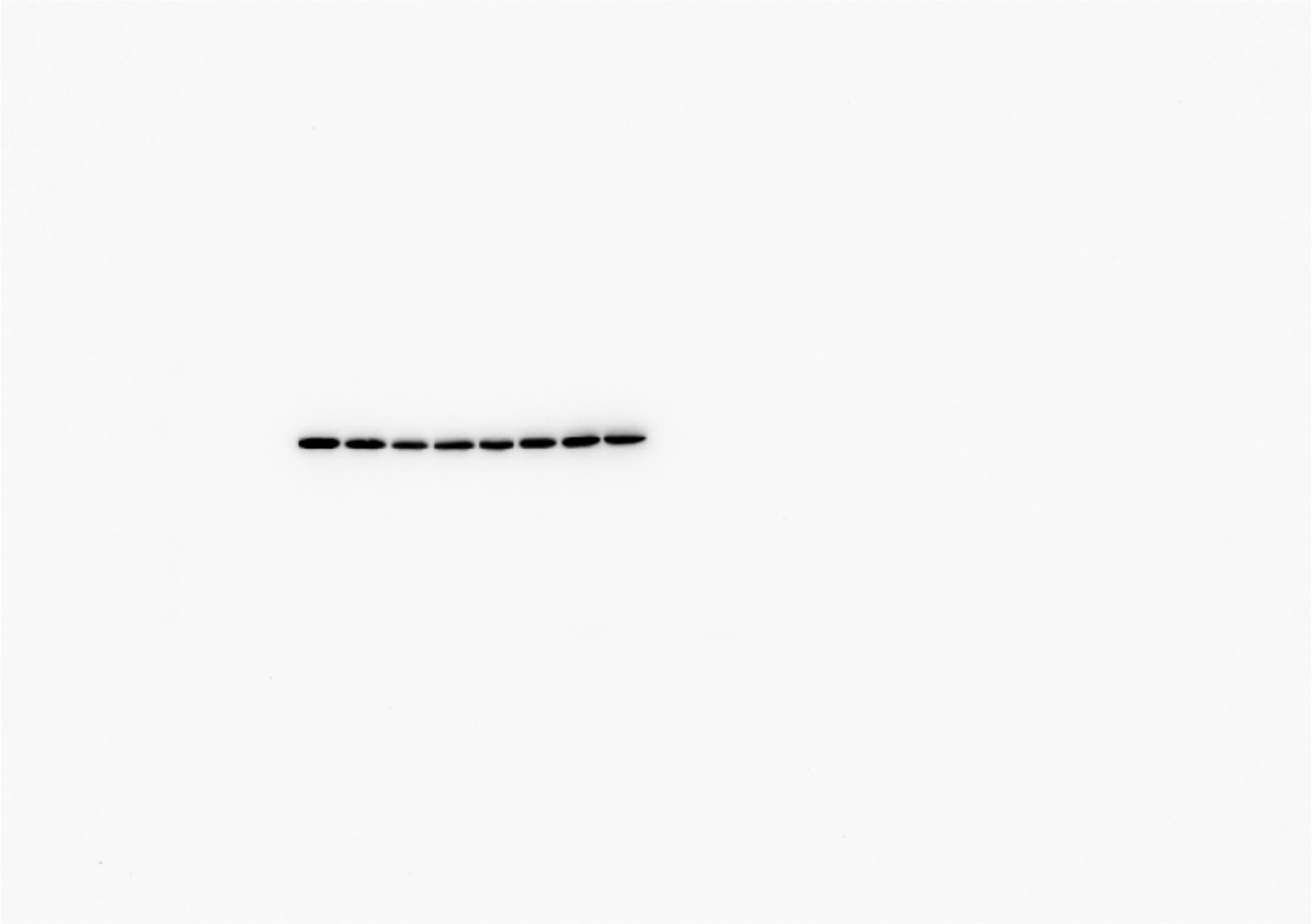

**Supplementary Figure 22.** Raw western blot image beta-actin in Fig. 3F.

| Analyte | WT Control (n=12) | WT DOCA (n=12) | IL17KO Control (n=5) | IL17KO DOCA (n=6) |
|---|---|---|---|---|
| BUN (mg/dL) | 25.00 ± 0.61 | 13.25 ± 1.00** | 27.1 ± 1.32 | 14.08 ± 1.91* |
| Creatinine (mg/dL) | 0.20 ± 0.01 | 0.19 ± 0.01 | 0.18 ± 0.01 | 0.18 ± 0.01 |
| BUN/CREA ratio | 130.89 ± 8.43 | 68.94 ± 3.17** | 142.54 ± 16.30 | 69.16 ± 14.25** |
| Total protein (g/dL) | 4.53 ± 0.14 | 4.97 ± 0.09* | 4.68 ± 0.23 | 5.01 ± 0.23 |
| Albumin (g/dL) | 2.64 ± 0.07 | 2.97 ± 0.06** | 2.76 ± 0.14 | 2.91 ± 0.15 |
| Globulin (g/dL) | 1.89 ± 0.07 | 2.00 ± 0.04 | 1.92 ± 0.11 | 2.10 ± 0.09 |
| A/G ratio | 1.40 ± 0.02 | 1.49 ± 0.03 | 1.46 ± 0.09 | 1.40 ± 0.04 |
| P (mg/dL) | 9.39 ± 0.52 | 8.56 ± 0.54 | 7.98 ± 0.35 | 7.72 ± 0.50 |
| Ca (mg/dL) | 8.49 ± 0.14 | 8.76 ± 0.17 | 9.10 ± 0.36 | 9.38 ± 0.32 |
| Na (mEq/L) | 146.06 ± 0.38 | 154.75 ± 1.05* | 147.00 ± 1.53 | 163.00 ± 4.78** |
| K (mEq/L) | 8.74 ± 0.45 | 6.01 ± 0.60 | 7.07 ± 0.28 | 5.58 ± 0.23 |
| Cl (mEq/L) | 110.75 ± 0.98 | 106.42 ± 1.14* | 122.00 ± 4.16 | 115.25 ± 2.18 |
| Na/K Ratio | 17.38 ± 0.84 | 28.17 ± 2.35** | 22.33 ± 0.33 | 27.5 ± 2.90 |

**Supplementary Table 1.** Serum metabolic panel at 21 days of control or DOCA-salt in wild-type (WT) and IL17-deficient (IL17KO) mice.
Data are presented as mean ± SEM. *p<0.05, **p<0.01 vs control, intergroup differences analyzed by two-way ANOVA with Tukey's multiple comparison test.

| Group | | IL-17 (pg/ml) | N |
|---|---|---|---|
| IL17KO | | | |
| WT | Control | 0.920 ± 0.1008 | 11 |
| | DOCA | 1.928 ± 0.2096* | 12 |
| IL17KO | Control | n.d. | 10 |
| | DOCA | n.d. | 10 |
| AAV-BR1-iCre | | | |
| $WT^{BR1}$ | Control | 0.870 ± 0.1284 | 7 |
| | DOCA | 1.912 ± 0.1156* | 6 |
| $IL17RA^{bECKO}$ | Control | 0.613 ± 0.1565 | 7 |
| | DOCA | 1.713 ± 0.1869* | 6 |
| Clodronate (CLO) | | | |
| PBS | Control | 0.789 ± 0.1573 | 7 |
| | DOCA | 1.940 ± 0.2179* | 7 |
| CLO | Control | 0.747 ± 0.1332 | 7 |
| | DOCA | 1.930 ± 0.1440* | 7 |
| BM Chimeras | | | |
| WT→WT | Control | 0.768 ± 0.0860 | 8 |
| | DOCA | 1.958 ± 0.2094* | 9 |
| $IL17RA^{-/-}$→WT | Control | 0.990 ± 0.0708 | 8 |
| | DOCA | 1.988 ± 0.2159* | 9 |
| $Nox2^{-/-}$→WT | Control | 0.647 ± 0.1002 | 6 |
| | DOCA | 2.023 ± 0.3110* | 9 |
| $Mrc1^{CreERT2}$ | | | |
| $BAM^{WT}$ | Control | 0.657 ± 0.0431 | 2 |
| | DOCA | 1.824 ± 0.3902 | 4 |
| $BAM^{IL17RA-/-}$ | Control | 0.709 ± 0.2537 | 3 |
| | DOCA | 1.514 ± 0.4713 | 4 |
| FTY720 | | | |
| Vehicle | Control | 0.625 ± 0.1054 | 12 |
| | DOCA | 2.111 ± 0.2199* | 9 |
| FTY720 | Control | 0.709 ± 0.1306 | 9 |
| | DOCA | 1.885 ± 0.2221* | 12 |
| i.c.v. losartan | | | |
| Saline | Control | 0.839 ± 0.1429 | 3 |
| | DOCA | 2.190 ± 0.3958* | 4 |
| Losartan | Control | 0.773 ± 0.1301 | 3 |
| | DOCA | 2.119 ± 0.1666* | 5 |

**Supplementary Table 2.** Serum IL-17 at 21 days of DOCA-salt.
Data are presented as mean ± SEM. *p<0.05 vs respective control, analyzed by unpaired two-tailed t-test for WT control vs DOCA, and two-way ANOVA with Bonferroni's multiple comparison test for all other groups.

| Target | Clone | Conjugate | Concentration | Catalog | Supplier |
|---|---|---|---|---|---|
| FACS | | | (ng per stain) | | |
| CD3e | 145-2C11 | APC/Cy7 | 25 | 100330 | Biolegend |
| CD4 | RM4-5 | PE/Cy5 | 100 | 100514 | Biolegend |
| CD8a | 53-6.7 | AF700 | 100 | 100730 | Biolegend |
| CD11b | M1/70 | AF700 | 8 | 101222 | Biolegend |
| CD11b | M1/70 | APC/Cy7 | 8 | 101226 | Biolegend |
| CD19 | 6D5 | APC/Cy7 | 25 | 115530 | Biolegend |
| CD36 | MF3 | FITC | 500 | MA5-16832 | Invitrogen |
| CD36 | MF3 | PE | 500 | MA5-28168 | Invitrogen |
| CD45 | 30-F11 | APC | 30 | 103112 | Biolegend |
| CD45 | 30-F11 | BV510 | 30 | 103138 | Biolegend |
| CD206 (intracellular) | C068C2 | APC | 62.5 | 141707 | Biolegend |
| Ly-6C | HK1.4 | BV711 | 15 | 128037 | Biolegend |
| Ly-6G | 1A8 | APC/Cy7 | 50 | 127623 | Biolegend |
| NK1.1 | PK136 | APC/Cy7 | 25 | 108723 | Biolegend |
| Tcrβ | H57-597 | PE/Cy7 | 15 | 109222 | Biolegend |
| Tcrβ | H57-597 | APC/Cy7 | 15 | 109220 | Biolegend |
| Tcrγδ | GL3 | APC | 15 | 118116 | Biolegend |
| Tcrγδ | GL3 | APC/Fire750 | 15 | 118135 | Biolegend |
| Immunofluorescence | | | (dilution) | | |
| CD31 | 2H8 | - | 1:100 | MAB1398Z | Sigma |
| CD31 | Polyclonal | - | 1:100 | AF3628 | R&D Systems |
| CD206 | MR5D3 | - | 1:200 | MCA2235 | Biorad |
| Claudin 11 | Polyclonal | - | 1:100 | 36-4500 | Invitrogen |
| Iba1 | Polyclonal | - | 1:100 | 019-19741 | Wako |
| GFP | Polyclonal | AF488 | 1:200 | A-21311 | Invitrogen |
| GFAP | Polyclonal | - | 1:100 | AB5804 | Millipore |
| Western Blotting | | | (dilution) | | |
| β-actin | AC-15 | - | 1:10,000 | A5441 | Sigma |
| eNOS | Polyclonal | - | 1:1,000 | 9572S | Cell Signaling |
| eNOS | Polyclonal | - | 1:1,000 | 07-520 | Millipore |
| pThr$^{495}$ eNOS | Polyclonal | - | 1:500 | 9574S | Cell Signaling |

**Supplementary Table 3.** Antibodies utilized for flow cytometry and flow activated cell sorting (FACS), immunofluorescence, and Western Blotting.

| Gene | | Primer sequence |
|---|---|---|
| mRNA primers | | |
| *Hprt* | m_Hprt_Real.1 | 5’-AGTGTTGGATACAGGCCAGAC-3’ |
| | m_Hprt_Real.2 | 5’-CGTGATTCAAATCCCTGAAGT-3’ |
| *Il17a* | m_Il17a_02_03_1 | 5’-CAGACTACCTCAACCGTTCCA-3’ |
| | m_Il17a_02_03_2 | 5’-AGAATTCATGTGGTGGTCCAG-3’ |
| *Agtr1a* | m_Agtr1a_Real_1 | 5’-AGAAAGCCATCACCAGATCAAG-3’ |
| | m_Agtr1a_Real_2 | 5’-GGGGCAGTCATCTTGAATTCTT-3’ |
| *Ren1* | m_Ren1_05_06_3 | 5’-GGGTGCTAAAGGAGGAAGTGT-3’ |
| | m_Ren1_05_06_4 | 5’-AGGAGTCAGTCTTGCTGATGC-3’ |
| *Gfap* | m_Gfap_Real_1 | 5’-ATTCGCACTCAATACGAGGCA-3’ |
| | m_Gfap_Real_2 | 5’-AGGTCTGCAAACTTAGACCGA-3’ |
| *Aldh1a1* | m_Aldh1a1_Real_1 | 5’-ACTGCTATATGATGTTGTCAGCC-3’ |
| | m_Aldh1a1_Real_2 | 5’-AGACCATGTTCACCCAGTTCTC-3’ |
| *P2ry12* | m_P2ry12_Real_1 | 5’-CAACAGATGCCAGTCTGCAAG-3’ |
| | m_P2ry12_Real_2 | 5’-ACATCCATGGTCCTGGTTCTG-3’ |
| *Trem2* | m_Trem2_Real_1 | 5’-ACTTCAGATCCTCACTGGACC-3’ |
| | m_Trem2_Real_2 | 5’-CTCCTGGCTGGACTTAAGCTG-3’ |
| *Tmem119* | m_Tmem119_Real_1 | 5’-ACCCAGAGCTGGTTCCATAG-3’ |
| | m_Tmem119_Real_2 | 5’-GAGTGACACAGAGTAGGCCA-3’ |
| *Cx3cr1* | m_Cx3cr1_Real_1 | 5’-CCATCTGCTCAGGACCTCAC-3’ |
| | m_Cx3cr1_Real_2 | 5’-GGTTCCAAAGGCCACAATGTC-3’ |
| *Cx3cl1* | m_Cx3cl1_03_03_1 | 5’-TCCCATAGCATTCTCCAAGAG-3’ |
| | m_Cx3cl1_03_03_2 | 5’-TCCTTGCTTCAGCAGTCACTT-3’ |
| *Cxcl12* | m_Cxcl12_1 | 5’-GTCTAAGCAGCGATGGGTTC-3’ |
| | m_Cxcl12_2 | 5’-TAGGAAGCTGCCTTCTCCTG-3’ |
| *Il6* | m_IL6_real.1 | 5’-ATGGATGCTACCAAACTGGAT-3’ |
| | m_IL6_real.2 | 5’-TGAAGGACTCTGGCTTTGTCT-3’ |
| *Il1b* | m_IL1b_05_06_1 | 5’-CTCTCCACCTCAATGGACAGA-3’ |
| | m_IL1b_05_06_2 | 5’-TTTTGTCGTTGCTTGGTTCTC-3’ |
| Genomic primers | | |
| *Icam* | m_ICAM1_prom.3 | 5’-GGACTCACCTGCTGGTCTCT-3’ |
| | m_ICAM1_prom.4 | 5’-GAACGAGGGCTTCGGTATTT-3’ |
| *Il17ra* exon 3 | m_Il17ra_exon3_Real_1 | 5’-AAAAACCTGACCCCGTCTTCC-3’ |
| | m_Il17ra_exon3_Real_2 | 5’-GGTCCACTCAACATGCAACAC-3’ |
| *Il17ra* exon 5 | m_Il17ra_exon5_Real_1 | 5’-TTCCTTCAGCCACTTTGTGGT-3’ |
| | m_Il17ra_exon5_Real_2 | 5’-CTTGGATTTGTGGTTTGGGTCC-3’ |

**Supplementary Table 4.** PCR primers
